# Reduced synaptic plasticity and E/I imbalance drive Peripersonal Space boundaries expansion in Schizophrenia

**DOI:** 10.1101/2024.07.21.604515

**Authors:** Renato Paredes, Vlad Grigoras, Francesca Ferroni, Martina Ardizzi, Francesca Ferri, Peggy Seriès

## Abstract

Abnormal encoding of peripersonal space (PPS) is believed to affect bodily self disruptions in schizophrenia (SCZ). Empirical studies show that SCZ patients exhibit a narrower PPS than controls but maintain its plasticity. Computational research links this smaller PPS to increased excitation of sensory neurons and reduced feedforward synaptic density. However, it is unclear how such differences influence learning during the expansion of PPS boundaries. We hypothesise that Hebbian plasticity can account for PPS expansion after active tool use training. To explore the effect of such mechanisms on PPS plasticity, we developed a SCZ network model which was fit to behavioural data before and after tool manipulation. We found that PPS expansion occurs in spite of E/I imbalance or reduced synaptic density, but does not match the post-training PPS representation of patients. A better fit was obtained after altering plasticity by either reducing the learning rate, increasing the forgetting rate or increasing the plasticity threshold. We discuss our findings in terms of dysfunctional plasticity in SCZ and highlight the key challenges in identifying the neurobiological correlates of reduced plasticity within PPS networks. Because current empirical data supports multiple viable mechanisms, we propose experiments to distinguish between the proposed plasticity accounts and clarify mixed findings on PPS representation in SCZ.

**Graphical Abstract:** 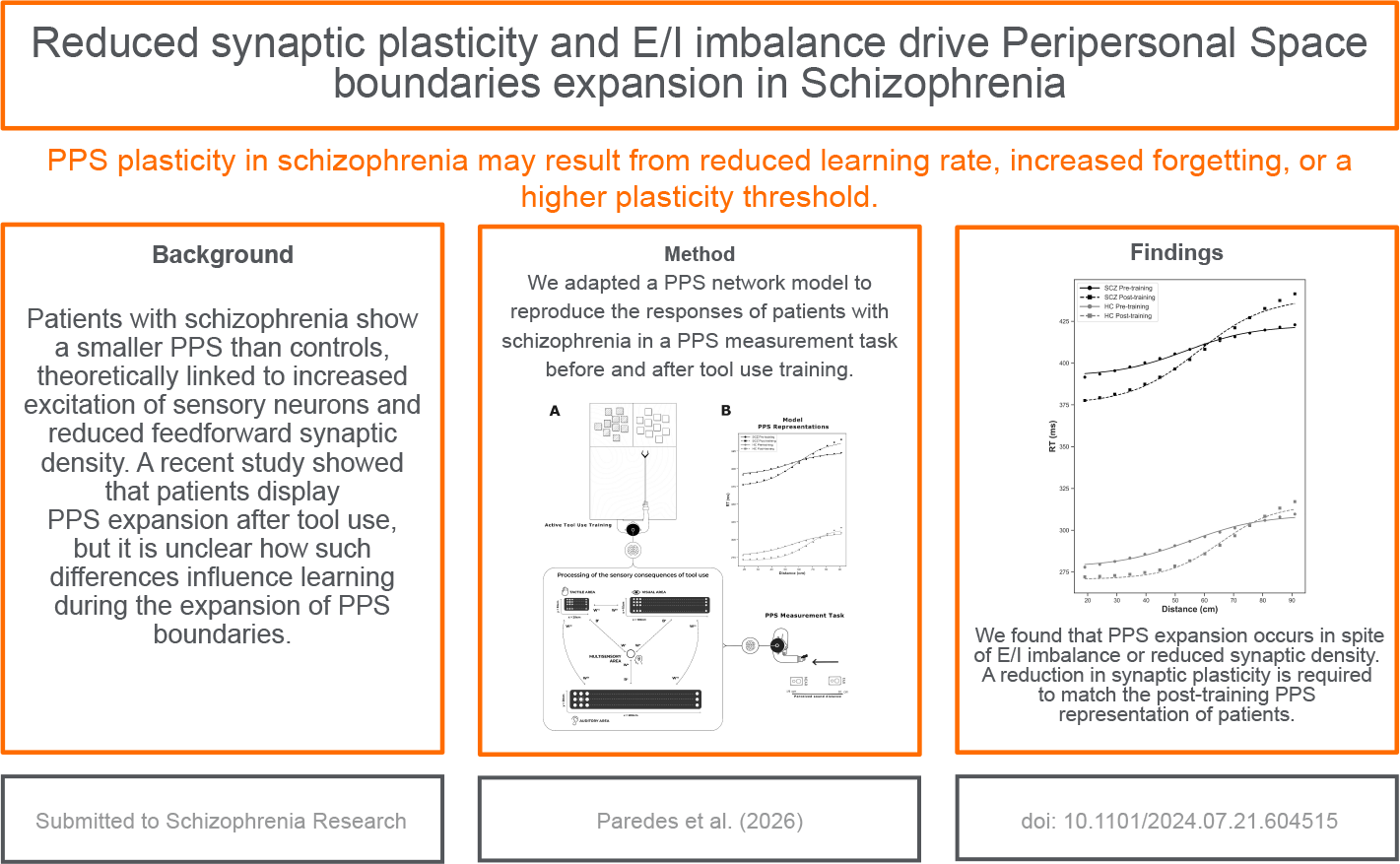

**Highlights:**

- Using a peripersonal space network model, we found that PPS expansion after tool-use occurs in spite of E/I imbalance or reduced synaptic density.
- Reduced feedforward synaptic plasticity is required to match the post-training PPS representation of patients.
- Such reduction of synaptic plasticity could be achieved by either reducing the learning rate, increasing the forgetting rate or increasing the plasticity threshold relative to a healthy control model.
- Our model predicts that measuring PPS at intermediate time points during a longer stimulation protocol would help distinguish between these plasticity differences.

## 1. Introduction

The brain encodes the space surrounding the body to facilitate interaction with the environment. Space is categorised based on proximity into peripersonal space (PPS), which is the space within arm’s reach, and extrapersonal space (EPS), which extends beyond that reach (van der Stoep et al., 2017). This classification derives support from research on fronto-parietal neurons in both macaques (Fogassi et al., 1996; Graziano et al., 1999) and humans (di Pellegrino et al., 1997; Làdavas et al., 1998; Gentile et al., 2013; Brozzoli et al., 2014; Ferri et al., 2015; Bernasconi et al., 2018; Noel et al., 2019), showing increased neural responses to visual and auditory stimuli occurring in close proximity to the body (refer to Grivaz et al. (2017) for a comprehensive review).

The PPS is not static, but constantly changing due to interaction with stimuli close to the body. This is orchestrated by PPS-encoding plastic, multimodal (auditory, tactile, and visual) fronto-parietal neurons that expand their receptive field to encompass the space around objects (tools) in contact with the body (Cléry et al., 2015), as observed in macaques (Iriki et al., 1996) and humans (Di Pellegrino and Làdavas, 2015). This plasticity has been observed in different training paradigms that involve tools (Magosso et al., 2010a), concurrent multimodal stimuli (Serino et al., 2015), echolocation (Tonelli et al., 2019), and social paradigms (Teneggi et al., 2013; Noel et al., 2022), with observed changes in the size and sharpness of the PPS boundary (Serino, 2019, 2022).

Patients with schizophrenia have narrower PPS boundaries compared to healthy controls (Di Cosmo et al., 2017; Lee et al., 2021; Ferroni et al., 2022; but see Noel et al., 2020). In terms of the sharpness of the boundary, the results are mixed, with some studies reporting steeper slopes than in controls (Di Cosmo et al., 2017) and others shallower (Ferroni et al., 2022) or not different (Noel et al., 2020; Lee et al., 2021).

We previously suggested (Paredes et al., 2022) that the reduction in PPS size reflects either increased excitation of unisensory neurons encoding stimulus location or reduced synaptic density between unisensory and multisensory neurons, with the latter also leading to a sharper slope. Increased excitation has also been theoretically linked to a decrease in tactile spatiotemporal discrimination (Lenzenweger, 2000; Chang and Lenzenweger, 2001, 2005; Michael and Park, 2016; Costantini et al., 2020; Ferri et al., 2016) and the tendency to experience self-aberrations (Michael and Park, 2016; Thakkar et al., 2011; Costantini et al., 2020).

Following this view and evidence of multisensory (Ferri et al., 2017; Di Cosmo et al., 2021) and sensorimotor (Kaufmann et al., 2015; Rossetti et al., 2020; Hirjak et al., 2021) integration abnormalities in schizophrenia, one would expect these differences to impact how PPS changes after motor training using tool manipulation, compared to what is observed in controls. An early study showed a reduced expansion of PPS (i.e., PPS size after training minus PPS size before training) after tool use in healthy individuals with high schizotypal traits (Ferroni et al., 2020). However, a follow up study did not observe differences in PPS expansion between individuals with SCZ and healthy controls after tool use (Ferroni et al., 2022), with patients exhibiting steeper PPS boundaries compared to before tool use (Ferroni et al., 2022). More recently, it has been found that within SCZ individuals, those with higher autistic traits or early onset of the disease show sharper PPS boundaries after training (Lucarini et al., 2025).

The neural basis underlying the plasticity of PPS observed in patients with schizophrenia is unknown (Ferroni et al., 2022). A comprehensive explanation of how neural plasticity is influenced by E/I imbalance and reduced synaptic density in the network encoding for PPS is lacking. As yet it is unclear how learning mechanisms work in SCZ during PPS expansion. Moreover, there is no clear understanding of how plasticity of PPS in SCZ is related to the motor and multisensory integration differences observed in the schizophrenia spectrum (Bédard et al., 2000; Pedersen et al., 2008; Schwartz et al., 2003; Ferri et al., 2017; Di Cosmo et al., 2021).

The current study aims at implementing a computational model that can account for PPS plasticity in SCZ (Ferroni et al., 2022) and is compatible with the neurobiological mechanisms (hypofunction of NMDA receptors or increased dopaminergic activity) proposed by state-of-the-art computational modelling of psychosis (Friston et al., 2016; Sterzer et al., 2018; Lanillos et al., 2020; Jeganathan and Breakspear, 2021). A neural network model of PPS in SCZ (Paredes et al., 2022) is adapted to reproduce the expansion of PPS after a simulated audio-visuo-tactile training routine. The simulated results are compared to the experimental data of patients with schizophrenia before and after tool use (Ferroni et al., 2022).

## 2. Methods

### 2.1. Participants

We modelled a subset of the data from Ferroni et al. (2022), originally comprising 59 individuals, 27 with SCZ and 32 healthy controls (HC). We included only participants whose PPS representation before and after training was well characterised by a sigmoidal curve (see Section 2.2), excluding 17 participants (6 HC, 11 SCZ) with poor sigmoidal fits (*R*^2^ < .25). This sub-selection of participants (see Table 1) does not alter the findings on PPS plasticity reported in Ferroni et al. (2022).

**Table 1:** Demographic Information for SCZ and HC Groups and Clinical Scales of SCZ.

|  | SCZ | HC |
| --- | --- | --- |
| *n* | 16 | 26 |
| Male, *n* | 12 | 10 |
| Age, *years* (mean; SE) | 30.38; 2.68 | 27.27; 0.45 |
| Illness duration, *years* (mean; SE) | 6.73; 2.34 | n.a. |
| Chlorpromazine equivalent dose, *mg/die* (mean; SE) | 626.07; 93.21 | n.a. |
| **Scale and Subscales** |  |  |
| PANSS (mean; SE) |  |  |
| Total | 78.63; 4.08 | n.a. |
| Positive scale | 15.56; 1.45 |  |
| Negative scale | 21.38; 1.32 |  |
| General psychopathology scale | 41.69; 2.36 |  |

### 2.2. Peripersonal space task

The computational model simulates the experiment conducted on patients with schizophrenia described in Ferroni et al. (2022). Participants were exposed to 3000 ms auditory stimuli, increasing from 55 to 70 dB, and tactile stimulation on the right hand at intervals between 300 and 2700 ms from the sound onset (for comprehensive details, refer to Ferroni et al. (2022)). Changing the timing of tactile stimulus affects spatial perception, with longer delays from sound onset making the sound seem closer to the hand. Participants were told that the auditory stimulus was unrelated to the task and had to press a button as quickly as possible in response to tactile stimulation. Faster reactions to the tactile stimulus are observed when the sound is perceived to be near the body (see Figure 1).

**Figure 1:**
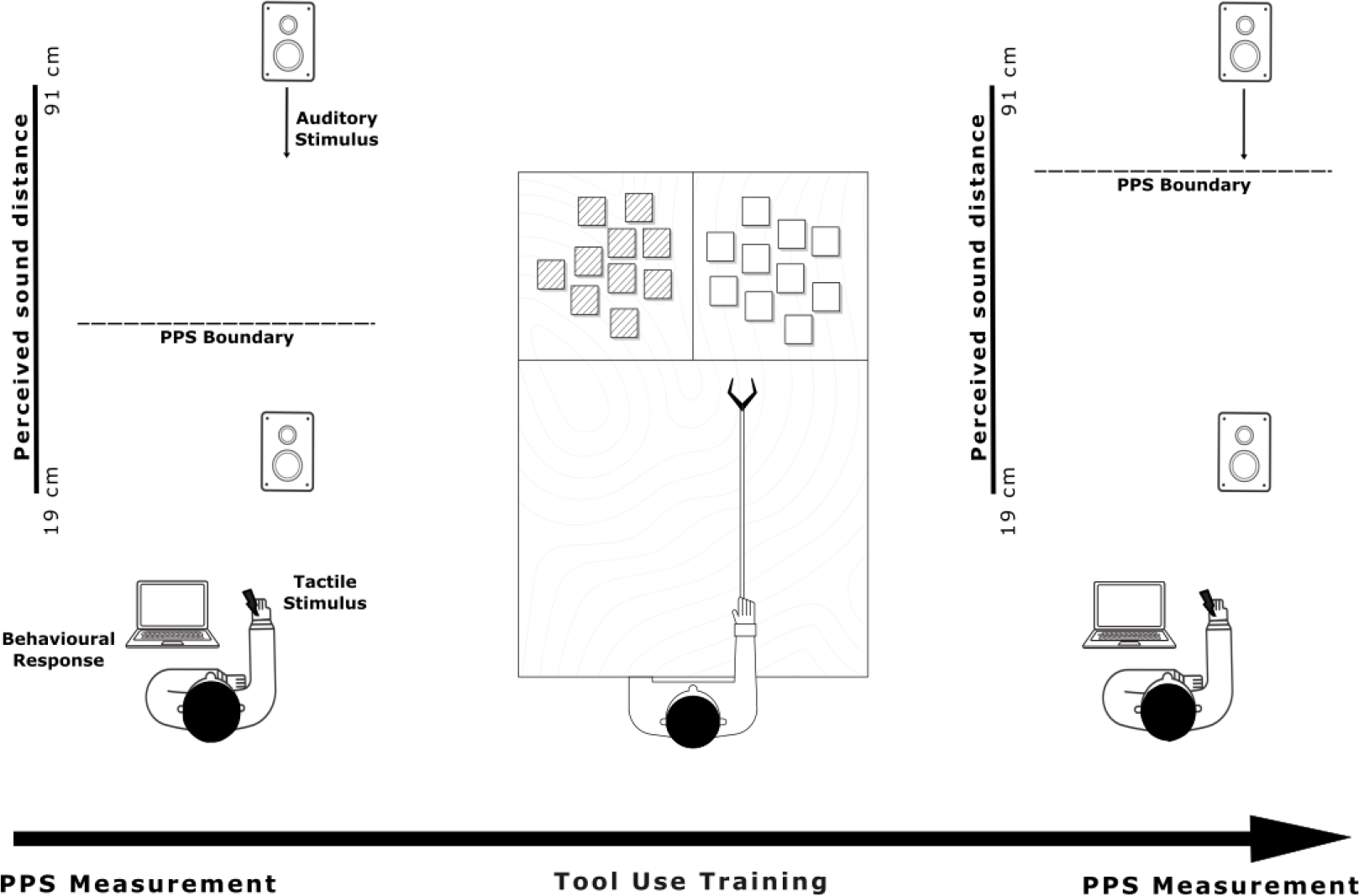
Graphical representation of the PPS measurement ask and the active tool use training protocol. The figure illustrates the experimental sequence adopted for participants in the original study Ferroni et al. (2022): measurement of PPS representation (left and right, Section 2.2) conducted both prior to and following the active tool-use training phase (centre, Section 2.3). This experimental manipulation leads to an expansion of the boundaries of the PPS.

Reaction times (RTs) averaged for each participant across different delay intervals were fitted by a sigmoidal function. The resultant sigmoidal curve is interpreted as the PPS representation of each participant (Magosso et al., 2010a,b; Di Cosmo et al., 2017; Lee et al., 2021). Quick RTs are interpreted as corresponding to sensory regions within the PPS, while slow RTs point to regions beyond the PPS. As is common in this literature, we interpret the central point (CP) of the curve as the size of the PPS, and the slope as the steepness of the PPS boundary. We characterised the group-level responses before and after tool use using a sigmoidal function parameterised by the median CP and median slope estimated for each group. We deviated from the Spearman-Karber method used in the original study (Ferroni et al., 2022) because our network model inherently holds the assumption of a sigmoidal curve as the underlying psychometric function. As previously reported (Ferroni et al., 2022), when using this method, the differences in the PPS boundary observed in the SCZ group before and after tool use are no longer evident. Therefore, we will not seek to reproduce them with our model.

### 2.3. Active tool use training

The training consisted in using a tool to reach objects beyond arm reach, a method widely recognized for expanding the expansion of the boundaries of the PPS representation (Iriki et al., 1996; Di Pellegrino and Làdavas, 2015). Participants were instructed to move 50 objects in their far space using a 75 cm garbage clamp (60 cm aluminium shaft, 12 cm lever) (see Figure 1).

The participants move the objects along the short side of a table and then back, for a total of 100 movements and a typical duration of 10 minutes. This protocol (Ferroni et al., 2020) is based on empirical evidence showing that the use of the tool extends the boundaries of the PPS representation by making out-of-reach objects ‘reachable’ (Bassolino et al., 2010; Canzoneri et al., 2013), which is indexed by the expansion of the visual receptive fields of intraparietal sulcus neurons towards the part of the space where the tool operates (Iriki et al., 1996).

### 2.4. An updated neural network model of PPS representation

We adapted the neural network employed by Paredes et al. (2022) to model the sensory processing of healthy controls and patients with schizophrenia in the two tasks conducted in the study (Ferroni et al., 2022): PPS representation measurement through audio-tactile stimuli, and sensory consequences of tool use training (See Figure 2).

**Figure 2:**
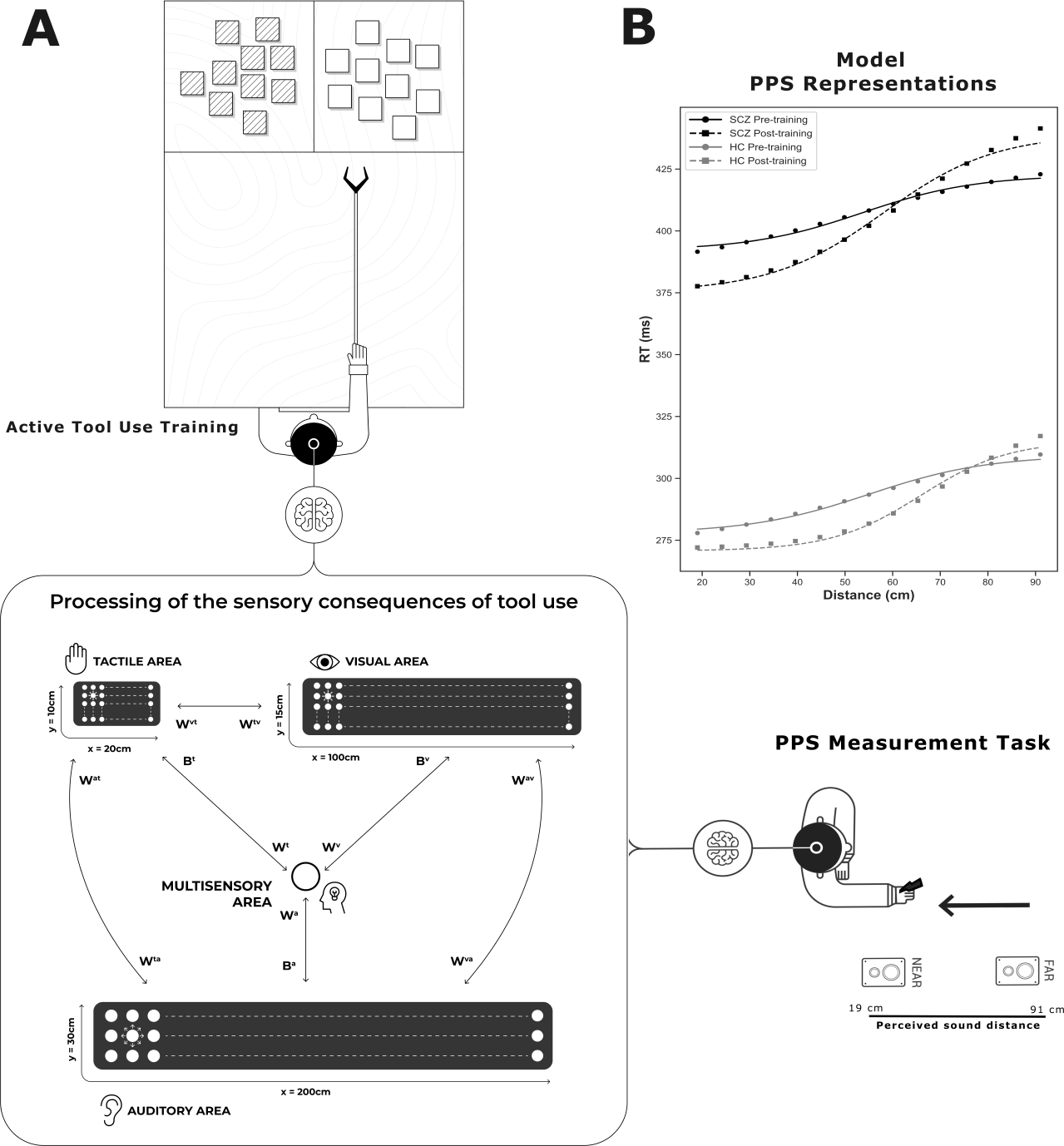
Model of the expansion of the PPS audio-visuo-tactile representation generated by the processing of the sensory consequences of tool use training. **Panel A** shows the network model (bottom left, Section 2.4), which is composed of three unisensory areas (tactile, auditory, and visual) connected to a multisensory area. The unisensory regions are configured to encode the spatial dimensions of the hand (10 cm x 5 cm), the external auditory space (200 cm x 30 cm) and the visual field (100 cm x 15 cm), respectively. The unisensory areas are connected with each other through cross-modal synapses, and with the multisensory area through feedforward and feedback synapses. This network is used to model both the responses of participants in the PPS measurement task (bottom right, Section 2.2) and the sensory consequences of motor activity during the active tool use training protocol (top left, Section 2.3). **Panel B** shows group averaged RTs (markers) of participants in the PPS measurement task and the PPS representations (lines) generated by the network models (Sections 2.5 and 2.6) before and after training. Our PPS expansion model was able to reproduce changes in the size of the PPS representation after active tool-use in HC and SCZ (top right, Sections 3.1 and 3.2).

The model describes three unisensory areas (auditory, visual, and tactile) connected with a multisensory area that contains multisensory representations of the PPS (see Figure 2 for an illustration). Neurons in unisensory areas are characterised by their receptive field (RF) defined in hand-centred coordinates (along the horizontal and vertical axis) and respond to stimulation at specific spatial coordinates in relation to the hand. Sensory areas are modelled with different spatial resolutions (number of neurons per space covered) reflecting sensory modality spatial sensitivity: higher in tactile and visual (Duncan and Boynton, 2007) than auditory (Mills, 1958). The tactile area has 200 neurons in a *M*^*t*^ = 20 × *N*^*t*^ = 10 grid encoding a 20 cm × 10 cm skin region of the left hand; the auditory area has 60 neurons in a *M*^*a*^ = 20 × *N*^*a*^ = 3 grid covering a 200 cm × 30 cm auditory space on and around the hand; and the visual area has 1500 neurons in a *M*^*v*^ = 100 × *N*^*v*^ = 15 grid covering a 100 cm × 15 cm visual space on and around the hand.

All neurons in each area interact via lateral synapses according to a Mexican-Hat pattern (i.e., near-excitation and far-inhibition). Synaptic weights are derived from the difference between two Gaussian functions (excitatory and inhibitory), defined by parameters 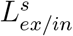 and 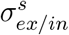, indicating curve height and width in each sensory modality *s*. Furthermore, sensory areas are linked by cross-modal synapses (*W*^*ss*^), where neurons from one sensory modality selectively excite neurons representing the corresponding spatial region in the other two modalities.

For simplicity, the multisensory area is composed of a single neuron connected to all neurons in the auditory, visual, and tactile areas, both through feedforward (*W*^*s*^) and feedback (*B*^*s*^) synapses. Synaptic weights from/to the neurons that encode the near space of the hand (i.e., the PPS, delimited by the parameter *Lim*) are defined by the parameters 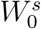 and 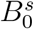, respectively. The feedforward and feedback synaptic weights exhibit an exponential decay proportional to the distance from *Lim*. This means that sensory inputs near the hand strongly influence the multisensory neuron, which in turn more strongly affects neurons encoding space near the hand than those encoding distant space. This configuration leads to multisensory neuron responses that align with the responses of neurons in parietal, temporo-parietal, and premotor regions that react to tactile stimuli on the body and to auditory stimuli presented close to, but not far from the body (Graziano et al., 1999; Schlack et al., 2005; Grivaz et al., 2017; Bernasconi et al., 2018). Model details are in the Supplementary Material.

### 2.5. PPS task simulation in the network

Following Paredes et al. (2022), the experimental task was simulated by making the following assumptions: First, the apparent velocity of the sound was 30 cm/s. Second, model simulations evaluated RTs for tactile stimuli presented with 15 different delays from the sound onset ranging from 300 to 2700 ms, which according to our first assumption correspond to 15 subjectively perceived sound distances between 19 cm and 91 cm from the hand. The RT of the network was defined as the time at which any neuron in the tactile area reached 90% of its maximum activation state. Third, a linear regression is applied to the raw RTs of the network to match the RTs of human participants, given that the model does not capture the entire neural process that causes the motor responses examined in the task. Fourth, to calibrate the network to reproduce PPS representations of healthy individuals, we fitted the parameters governing feedforward and feedback auditory synaptic weights outside the near-space of the hand to generate RTs that match the average PPS representation observed in the HC group before tool use (Ferroni et al., 2022) (see Figure 2B).

### 2.6. Training simulation in the network

To recreate the active tool use training in the model, we do not explicitly model motor processing and motor neurons activity but only the sensory consequences of each object movement. We assume that each object movement produces tactile (tool grip), visual (object position), and auditory (sound from interactions between the metal tool, the object, and the table) signals that we represent in the network as a synchronous audio-visuo-tactile stimulus.

The audio-visuo-tactile stimulus used to represent the sensory consequences of object movements is described by a 2D Gaussian function, where 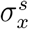 and 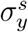 represent the horizontal and vertical spread of the stimulus, respectively (where *s* is either tactile, auditory or visual). The tactile stimulus is always delivered at the central point of the tactile area. Auditory and visual stimuli were applied at horizontal position 60 cm, centred vertically, to simulate the sensory effects of moving objects away at a distance equivalent to the shaft length.

Following Serino et al. (2015), we assume that the synchronicity between tactile stimulation in the hand and auditory or visual stimulation in the far space triggers the expansion of PPS via Hebbian Learning. The change in the synaptic weight, 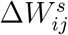, at each time step during the training simulation is defined as the result of a reinforcing factor and a forgetting factor. In the reinforcing factor the activity rates of the pre-synaptic and multisensory neurons are multiplied by the learning rate 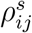 (*t*) (where *s* is either auditory and gradually reduces as the or visual^1^). The rate starts at a fixed level 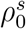 connection strengthens, reaching zero when the weight is at its maximum. Here the multisensory neuron activity is weighted against a small plasticity threshold *θ* to avoid strengthening in cases of residual activity. The forgetting factor is computed when the pre-synaptic neurons are inactive and the post-synaptic neuron has activity above threshold, leading to the decrease of the synaptic weights by the forgetting rate *κ*^*s*^.

The audio-visuo-tactile stimulus was presented 10 times to train the network. To emulate the expansion and slight sharpening of the PPS boundaries (Δ_*CP*_ = 15.84 cm, Δ_*Slope*_ = .042) observed in the HC group after active tool-use, we fitted the parameters that define the network learning rate 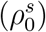 and the horizontal spread 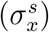 of the audiovisual stimulus, ensuring that the RTs of the HC model after training match the post-training PPS representation of the HC group (Ferroni et al., 2022). The fitted training procedure achieved the desired effect both in the size and in the slope of the PPS representation (Δ_*CP*_ = 10.80 cm, Δ_*Slope*_ = .027, see Figure 2B). More details of training are in the Supplementary Material.

### 2.7. Modelling the influence of SCZ in the network

The setup of the HC model was considered as a starting point to introduce alterations such as those observed in the schizophrenia spectrum (Di Cosmo et al., 2017; Lee et al., 2021; Noel et al., 2020; Ferroni et al., 2022). Following Paredes et al. (2022), we explored three putative mechanisms that could explain the pre-training PPS representation in the SCZ spectrum: excitation/inhibition (E/I) imbalance (Jardri et al., 2016), failures in top-down signalling (Sterzer et al., 2018), and decrease in synaptic density^2^ (Berdenis van Berlekom et al., 2019). We simulated the impact of these differences on the PPS representation to establish a baseline before examining the effects of active tool use training in SCZ.

The modulation of the E/I balance was implemented by increasing or decreasing the strength of the recurrent excitatory connectivity in each sensory area. This was modelled by increasing or decreasing the value of the parameter 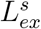 in all unisensory areas, as in our previous implementations (Paredes et al., 2022, 2025). The failures in top-down signalling were implemented by uniformly weakening top-down synaptic weights between multisensory and unisensory areas. This was achieved by changing the value of the parameter 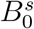 that governs the strength of the feedback synapses in all sensory areas.

In line with Paredes et al. (2022), the decrease in synaptic density between multisensory and unisensory connections was implemented by resetting the connection weights of feedforward synapses (*W*^*s*^) that were below a certain threshold *ρ* to zero. The same procedure was implemented to decrease the density of cross-modal synapses (*W*^*ss*^). The modelling details are in the Supplementary Material.

## 3. Results

### 3.1. Accounting for PPS differences in SCZ before training

To select the model that provides the best fit, we considered the Root Mean Square Error (RMSE) adjusted by the number of free parameters, as in Paredes et al. (2022, 2025). The results are compatible with our previous findings (Paredes et al., 2022): the network requires both enhanced recurrent excitation (*L*_*ex*_ = 0.79) within the unisensory areas and feedforward synaptic pruning (*ρW*^*s*^ = 0.48) to generate PPS representations that align with the responses observed in SCZ (adj RMSE = 0.35 ms), even with data from a different patient group (Ferroni et al., 2022). The optimal fit of this model to the SCZ group data before training is shown in Figure 2B. However, with the present data, we cannot discard a model based only on recurrent excitation (*L*_*ex*_ = 0.79, adj RMSE = 0.33 ms) and a model based only on feedforward pruning (*ρW*^*s*^ = 0.28, adj RMSE = 0.41 ms) to account for the PPS of this group of patients. The three models are empirically indistinguishable and achieve a goodness-of-fit equivalent to that observed in the HC model (adj RMSE = 0.21 ms).

### 3.2. Accounting for PPS after training in SCZ

We applied the training routine to our SCZ models to explore the resulting changes in the size and slope of the PPS representation (see Figure 3). These models broadly capture how tool use affects PPS representation in SCZ: expanding PPS without impacting its slope. However, none of the resulting PPS representations match the post-training SCZ data (see Table 2). We observe that lateral excitation enhances the effects of training in PPS expansion, whilst barely affecting its slope. In contrast, we do not observe the enhanced expansion effect in the feedforward pruning model. When both differences are present, the resulting post-training PPS representation is equivalent to that obtained with the lateral excitation model.

**Table 2:** Effects of tool use on PPS representation observed in SCZ and in the evaluated models. Δ represents the difference between post-training and pre-training.

| | $\Delta_{CP}$ | $\Delta_{Slope}$ | Adj RMSE |
| --- | --- | --- | --- |
| SCZ - $L_{ex}^s$ | 11.63 cm | .018 | 4.36 ms |
| SCZ - $\rho_{W^s}$ | 10.84 cm | .028 | 4.37 ms |
| SCZ - $L_{ex}^s$ - $\rho_{W^s}$ | 11.63 cm | .018 | 4.55 ms |
| SCZ - $L_{ex}^s$ - $\rho_{W^s}$ - $\rho_0^s$ | 4.45 cm | .000 | 1.67 ms |
| SCZ - $L_{ex}^s$ - $\rho_{W^s}$ - $\kappa^s$ | 4.45 cm | .000 | 1.67 ms |
| SCZ - $L_{ex}^s$ - $\rho_{W^s}$ - $\theta$ | 4.45 cm | .000 | 1.67 ms |

**Figure 3:**
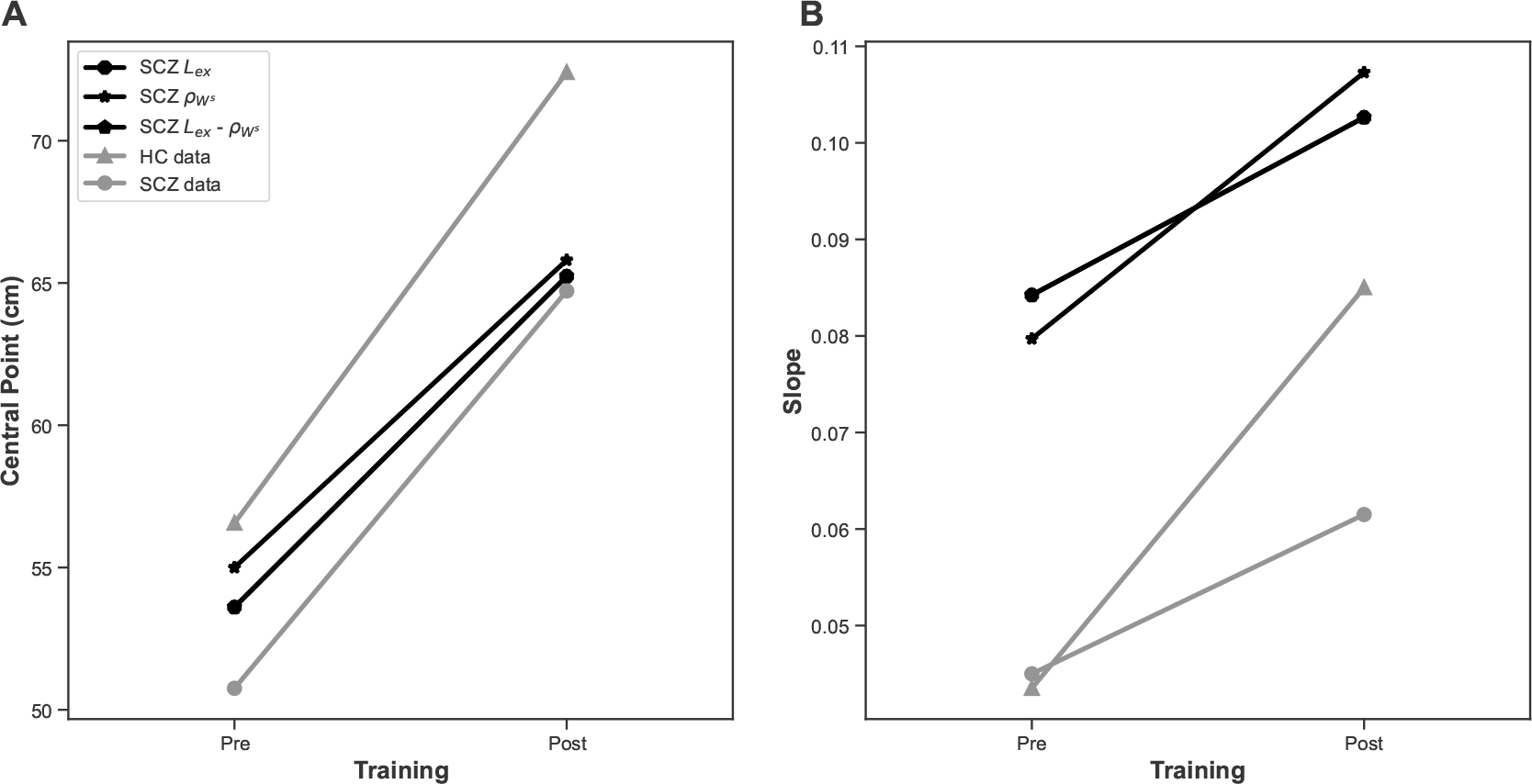
Effects of training in PPS representation in network models with SCZ differences. **Panels A** and **B** show the changes in PPS size and slope respectively. The black markers represent the data generated by the SCZ models in the simulated PPS task. The gray markers represent the data collected in the experiment (Section 2.2). The SCZ models display a larger PPS representation after training compared to before training. Note that the markers for the 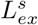 and 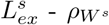 models overlap, as both models show equivalent size and slope before and after training.

The observed mismatch with empirical data suggests that the training effect should be subtler in the SCZ network. We examined the effect of fitting three parameters that govern feedforward synaptic plasticity: learning rate 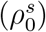, forgetting rate (*κ*^*s*^) and plasticity threshold (*θ*). We achieved an equivalent fit to the SCZ post-training data after reducing the learning rate (from 6.14 × 10^−2^ to 1.44 × 10^−2^), increasing the forgetting rate (from 5 × 10^−5^ to 1.25 × 10^−3^), or increasing the plasticity threshold (from .05 to .80) (see Table 2). All these models achieve the goal of expanding the PPS whilst keeping the slope constant and providing a good fit to the post-training SCZ data. Equivalent changes in the learning rule are necessary to reproduce the expansion effect observed in HC with the pre-training SCZ model (see Supplementary Material). The optimal fit of the 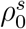 model to the SCZ group data after training is shown in Figure 2B.

To understand the impact of each plasticity parameter on changes in audio-tactile PPS representation, we explored the temporal evolution of feed-forward auditory weights (*W*^*a*^) during training in these candidate models (see Figure 4). Compared to baseline HC and SCZ models, we observe that training in these models has a weaker effect on increasing feedforward synaptic weights in neurons encoding the area outside the hand. Furthermore, we observe that weakening the reinforcing factor 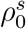 or *θ*) leads to a slower increase in weights and a smaller increase in PPS size during training. In contrast, strengthening the forgetting factor (*κ*^*s*^) leads to a faster increase in the weights that converges in fewer steps and a smaller initial PPS size increase followed by a plateau. We cannot distinguish between these models with current experimental data, but their distinct learning dynamics suggest a path for further empirical evaluation.

**Figure 4:**
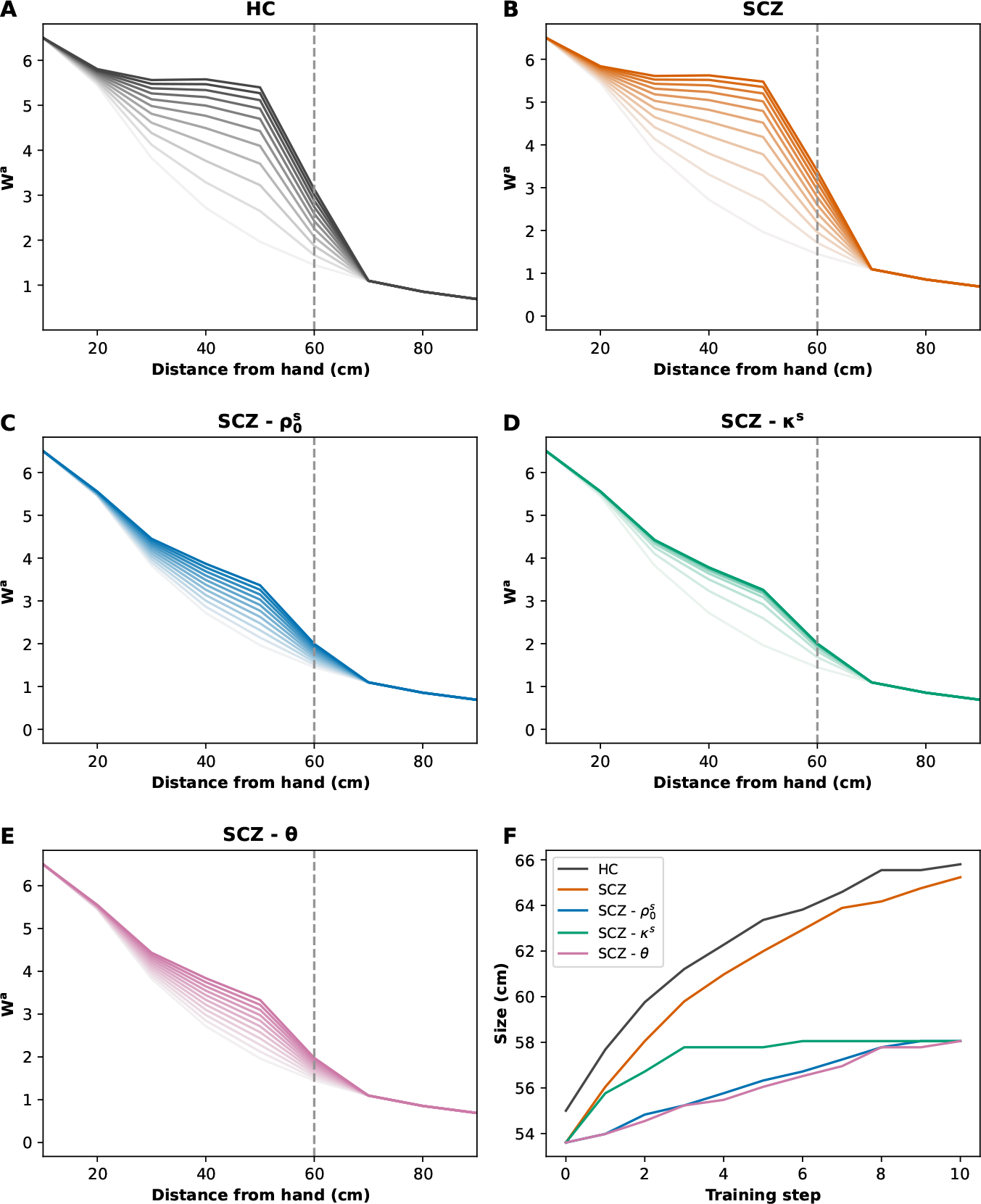
Dynamics of network models across training steps. **Panels A-E** show the feed-forward synaptic weights of the neurons encoding the auditory space (*W*^*a*^) as a function of the distance from the hand (*N*^*a*^ = 2). The gray dashed line represents the distance at which the audiovisual stimulus was presented. The colour-coded lines depict the evolution of the weights during a 10 step training, with lighter shades at the beginning and darker at the end. Compared to baseline HC and SCZ 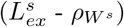 models, weakening the reinforcing factor 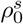 or *θ*) leads to slower synaptic plasticity, whereas strengthening the forgetting factor (*κ*^*s*^) leads to a quicker synaptic plasticity that converges in fewer steps. **Panel F** shows PPS size changes during training for each model. The figure reveals that compared to baseline models, weakening the reinforcing factor leads to a smaller and linear expansion in PPS size during training, whereas strengthening the forgetting factor leads to a smaller initial expansion followed by a plateau.

## 4. Discussion

This study aimed at modelling the neural mechanisms that give rise to the changes of PPS observed in SCZ after tool-use. For this purpose, an existing SCZ PPS network model (Paredes et al., 2022) was adapted to simulate audio-visuo-tactile stimulation under conditions of imbalance in E/I and decreased synaptic density to recreate the sensory consequences of active tool use that leads to the expansion of PPS in patients (Ferroni et al., 2022). These network conditions could reflect the consequences of the hypofunction of NMDA receptors, which characterise current computational accounts of SCZ (Friston et al., 2016; Sterzer et al., 2018; Lanillos et al., 2020).

Following Paredes et al. (2022), fitting to the experimental data (Section 3.1) suggests two possible neural mechanisms that account for the representations of PPS observed in SCZ patients before undergoing active tool use training: an increase in recurrent excitation of unisensory neurons or a decrease in feedforward synaptic density between unisensory and multisensory neurons. Both increased excitation and decreased synaptic density could explain the smaller size of PPS in patients (Di Cosmo et al., 2017; Lee et al., 2021; Ferroni et al., 2022), while a pronounced reduction of synaptic density could explain the increased slope steepness of PPS (Di Cosmo et al., 2017).

Assuming the existence of either mechanism, an additional assumption was required to reproduce the representation of PPS observed in SCZ patients after active tool use training (Section 3.2). The post-training PPS representation of patients could only be achieved after altering the feedforward plasticity of our SCZ model. This can be modelled by weakening the reinforcing factor or strengthening the forgetting factor within the Hebbian learning rule, producing equivalent outcomes but different dynamics (see Figure 4).

### 4.1. PPS plasticity in SCZ

A key distinction to address learning differences in SCZ (Friston et al., 2016; Sterzer et al., 2018; Guo et al., 2019) is whether impairments affect explicit learning (i.e. within conscious awareness) or implicit learning (i.e. outside of conscious awareness), given that both processes involve different neural mechanisms (Loonis et al., 2017). Active tool use training modelled in this report falls into the category of implicit learning (but refer to Bufacchi and Iannetti (2018) for a different perspective). The literature shows contradictory evidence about whether implicit learning is altered (Weickert, 2018; Haarsma et al., 2021; Tuominen et al., 2022) or intact (Barch et al., 2017; Orlov et al., 2021) in the schizophrenia spectrum. Notably, sensory learning alterations have been observed in patients with schizophrenia (Lev-Ari et al., 2015; Dzafic et al., 2021).

Our approach allowed us to examine neural plasticity in the context of an E/I imbalanced and pruned PPS network, suggesting the presence of reduced plasticity of auditory and visual feedforward synapses during an implicit learning task (see Section 3.2). This result is consistent with reinforcement learning studies involving SCZ patients showing a decreased or altered modulation of the learning rate in associative tasks, which are interpreted as plausible surrogates for alterations in synaptic plasticity (Diwadkar et al., 2008; Hernaus et al., 2018). Moreover, we found that, by itself, increased lateral excitation in sensory areas leads to the learning of an expanded new PPS representation out of concurrent audio-visuo-tactile stimulation (see Section 3.2). This observation broadly aligns with literature indicating that an elevated E/I ratio is usually accompanied with higher learning (van Bueren et al., 2023; Zacharopoulos et al., 2021).

Within our plasticity rule, the change in synaptic weights is proportional to neural activity. In the context of an imbalanced E/I network, increased excitation increases the plasticity of our model. Thus, during the fitting procedure the parameters of the Hebbian rule (learning rate, forgetting rate, or plasticity threshold) established for the HC network were adjusted to compensate for the hyper-plasticity of the pre-training SCZ network. This results in a hypo-plastic network that leads to a reduced increase in feedforward synaptic weights after training compared to the HC model (see Figure 4). Could this model behaviour be mapped onto accounts of dysfunctional plasticity in SCZ, where networks can fluctuate between transient or chronic hyper- and hypo-plastic states (Keshavan et al., 2015; Guterman et al., 2021)? To address this theoretical question, it is necessary to clarify whether the hypo-plasticity observed in the PPS network reflects the dysregulation of NMDARs by neurotransmitters such as dopamine, serotonin, or acetylcholine (Friston et al., 2016; Sterzer et al., 2018). Alternatively, it may instead index structural plasticity, a homeostatic mechanism in SCZ that progressively eliminates excitatory dendritic synapses in response to prolonged hyperplastic states (Palaniyappan, 2019). Another challenge is to clarify to what extent these neural mechanisms can be adequately represented by a Hebbian plasticity rule (Shouval et al., 2002; Cooper and Bear, 2012), or whether alternative approaches to PPS plasticity modelling are required (Bertoni et al., 2025; Bufacchi et al., 2025).

Current empirical evidence on PPS and its plasticity in SCZ has only begun to accumulate (Di Cosmo et al., 2017; Lee et al., 2021; Noel et al., 2020; Ferroni et al., 2020, 2022; Lucarini et al., 2025) and does not yet allow us to distinguish between the proposed plasticity differences. However, our models make predictions which, if tested experimentally in the future, could be used to this effect. Paradigms that measure PPS changes due to active tool use training or, more generally, synchronous sensory stimulation (Serino et al., 2015; Tonelli et al., 2019) could distinguish the proposed learning profiles in SCZ (weaker learning vs. stronger forgetting) if PPS is measured at intermediate time points between the onset and the end of the stimulation protocol (see Figure 4F). Further simulations shown in Supplementary Material suggest that PPS size will also vary after longer stimulation or when the stimulus is presented at different distances from the hand (e.g. with different shaft lengths). We believe that an in-depth examination of PPS learning dynamics may clarify the mixed findings on PPS representation (Di Cosmo et al., 2017; Lee et al., 2021; Noel et al., 2020) and plasticity (Ferroni et al., 2020, 2022; Lucarini et al., 2025) in SCZ.

### 4.2. Potential therapeutic venues for bodily self-aberrations

Unusual PPS representations have been theoretically linked to bodily self-aberrations in SCZ (Noel et al., 2017; Serino, 2019; Paredes et al., 2022). According to this view, weaker or unstable PPS representations observed in patients could be behind the higher proneness to experience bodily self-illusory phenomena, such as the observed in the Rubber Hand (Rossetti et al., 2020; Costantini et al., 2020; Zopf et al., 2021; He et al., 2022), Pinocchio (Michael and Park, 2016) and Enfacement (Ferroni et al., 2019; Sandsten et al., 2020) illusions. Presumably, this occurs because the multisensory integration of exteroceptive and proprioceptive signals within PPS is crucial for bodily self-inferences (Noel et al., 2018b,a; Chancel et al., 2021).

Empirical evidence showing PPS plasticity in SCZ (Ferroni et al., 2022; Lucarini et al., 2025) could open up the possibility of exploring treatments of bodily self-aberrations symptoms based on sensory (Machingura et al., 2018; Donde et al., 2019) or motor training protocols (Hirjak et al., 2021). Our modelling results suggest that the expansion of PPS occurs in the context of specific alterations of neural plasticity that are still slowing plasticity. Thus, potential sensorimotor therapeutic protocols may require pharmacological (Michalopoulou et al., 2015) or neurostimulation (Lefebvre et al., 2020; Narita et al., 2020) techniques aimed at modulating learning and E/I balance in multisensory integration networks.

### 4.3. Limitations and Future Directions

Our network model is obviously a gross simplification of reality, exhibiting several inherent limitations (see Paredes et al. (2022) for a full discussion). Our model was fit to group-averaged data. With larger datasets, future modelling efforts will benefit from individual-level fitting and cross-validation to evaluate the robustness of the proposed neural mechanisms.

Furthermore, our model does not consider motor processing, which is essential in tool use training (Bassolino et al., 2010; Canzoneri et al., 2013). The model assumes that the movements generated by the participants are the same and produce the same sequence of multisensory stimuli. As such, our model does not account for the differences that could occur when these multisensory stimuli are generated by active motion compared to passive conditions (Tonelli et al., 2019; Bufacchi and Iannetti, 2018). This is critical given the evidence of sensorimotor integration deficits observed in SCZ (Kaufmann et al., 2015; Rossetti et al., 2020; Hirjak et al., 2021). Moreover, at the neural level, the sensory and motor cortices are adjacent and closely interact (Flanders, 2011; Sohn et al., 2021).

Following Guterman et al. (2021), we also encourage the design of novel experimental paradigms that aim to measure long-term (hours to days) plasticity of PPS, as opposed to the short-term plasticity (seconds to minutes) induced by the training protocol employed in this study. This inquiry is crucial, as alterations in the long-term associative learning of statistical contingencies between self-generated actions and their afferent sensory outcomes could result in an impaired capacity to anticipate the afferent consequences of one’s own activities (Guterman et al., 2021). Consequently, this may lead to novel explanations of the role of PPS plasticity in delusions of alien control observed in SCZ (Serino, 2019).

## 5. Funding

F.F. is supported by the “Departments of Excellence 2023–2027” initiative of the Italian Ministry of Education, University and Research for the Department of Neuroscience, Imaging and Clinical Sciences (DNISC) of the University of Chieti-Pescara, and by the Italian Ministry of University and Research (MUR), funded by the European Union – NextGenerationEU, under the National Recovery and Resilience Plan (NRRP) CUP: D53D23020890001.

## 6. CRediT authorship contribution statement

**Renato Paredes:** Conceptualisation, Methodology, Software, Formal Analysis, Writing - Original Draft. **Vlad Grigoras:** Conceptualisation, Methodology, Software, Writing - Original Draft. **Francesca Ferroni:** Conceptualisation, Investigation, Writing - Review and Editing. **Martina Ardizzi:** Investigation, Writing - Review and Editing. **Francesca Ferri:** Conceptualisation, Investigation, Writing - Review and Editing. **Peggy Seriès:** Conceptualisation, Methodology, Supervision, Writing - Reviewing and Editing.

## 7. Declaration of competing interest

None.

## 8. Availability of data and materials

The code used to produce the simulations presented in this manuscript can be found at: https://github.com/renatoparedes/SCZ_PPS_Expansion_Model

## 9. Acknowledgements

The authors acknowledge Sara Cincă and Diego Herrera for their contribution in the graphics used in this paper.

## 1 Participants

We modelled the data gathered by Ferroni et al. (2022), involving a sample of 16 individuals diagnosed with schizophrenia (SCZ) alongside 26 healthy control participants (HC). Inclusion criteria for SCZ participants required: (1) a confirmed diagnosis of Schizophrenia as per DSM-5 guidelines; (2) an achieved stable clinical condition post-resolution of acute symptoms, characterized by a reduced severity in psychotic symptoms to low or mild intensity (Herz et al., 2002); and (3) provision of written informed consent. Schizophrenia diagnoses were substantiated using the DSM-5 Axis I criteria (SCID-5-CV) (First et al., 2015). All patients were administered a low to medium dose of a single atypical antipsychotic. Evaluation for the pharmacological treatment’s impact involved the conversion of the therapy into chlorpromazine equivalents, with no significant correlation to performance observed (Refer to Ferroni et al. (2022) Supplementary Materials). Participants were excluded based on: (1) any current mental disorder linked to a general medical condition or substance abuse; (2) cognitive impairments impacting test compliance; (3) sensory abnormalities in touch or hearing. For further information on this cohort, please consult Ferroni et al. (2022).

## 2 Peripersonal Space network model

The network model used in this study is an extension of previous models developed by Magosso and her collaborators (Serino et al., 2015; Magosso et al., 2010). In particular, we built on the implementation of the model that simulates the PPS representation of patients with SCZ made by our research group (Paredes et al., 2022). For this study, we implemented trimodal unisensory modelling, with cross-modal connections, and further synaptic links to a multi-sensory region.

### 2.1 Sensory Modalities and Inputs

The model is characterised by 3 unisensory modalities (visual, auditory and tactile) that are connected to a multisensory neuron through feedforward and feedback synapses.

The tactile region encodes a space of 10 x 20 cm, corresponding to the hand. The visual region encodes a space of 15 x 100 cm, mapping to the space around the hand and extending 80 cm ahead and 2.5 cm on each side. Similarly, the auditory region encodes a space of 30 x 200 cm, extending 10 cm to each side of the hand and 180 cm in the front. Within the same modality, proximal neurons facilitate lateral excitation amongst themselves, while inhibiting neurons that are more distal, resembling the characteristics of a Mexican Hat function. Across different modalities, neurons selectively provide excitatory influence to other neurons possessing receptive fields that are aligned with analogous spatial locations, facilitated by cross-modal connections.

Each neuron has its own receptive field, which is described according to the following Gaussian function:

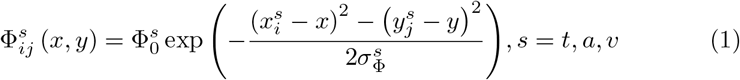

Here 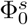 and 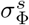 represent the amplitude and spread of the function, while *x & y* the spatial coordinates of the stimulus being applied. 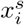 and 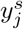 both represent the two dimensional RF centre of the neuron *i,j* from modality *s*. Given the large cortical area dedicated to the processing of stimuli falling on the hand (Longo and Haggard (2011)), tactile acuity will be the highest, with neurons extending 0.5 cms with respect to both Cartesian planes. The visual modality is slightly less sensitive, with neuronal RFs extending by 1 cm bidimensionally, whereas auditory neurons have large RFs, extending by 10 cms on both axes. Thus, RF centres are computed accordingly:

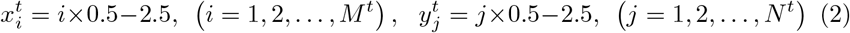

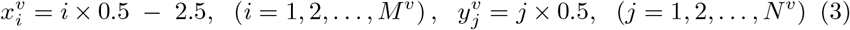

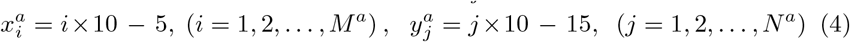

The receptive field of each neuron functions as its “focal point of awareness” for processing incoming stimuli. Neurologically, receptor cells act as transducers of light, pressure or vibration (depending on the sense) to electrical current. In the present model, the RF behaves as the intermediary between the stimuli and the network’s activity. The neuronal input is then computed as the convolution of the stimulus itself with the network’s receptive fields over the entire space, as such:

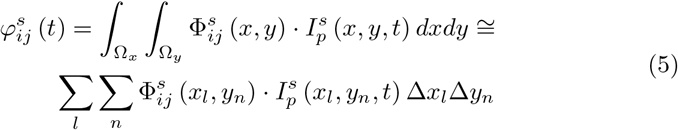

Here, Δ*x*_*l*_ = Δ*y*_*n*_ = 0.2cm is the discretising value over the coordinates space. 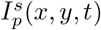 is the external stimulus applied to the tactile (*s* = *t*), auditory (*s* = *a*) or visual (*s* = *v*) modality, during training (*p* = *t*) or testing (*p* = *e*), at coordinates (x,y) and time step *t*.

### 2.2 Lateral Connections

All neurons in each unisensory modality interact via lateral synapses according to a Mexican-Hat function (i.e., near-excitation and far-inhibition). This is realised through the following Gaussian blob, where *ij* represents the post-synaptic neuron and *hk* the pre-synaptic one:

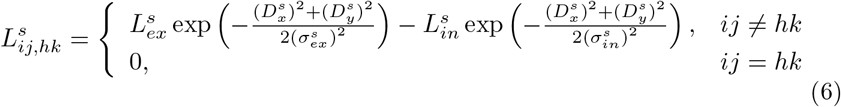

Here, 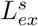 and 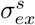 are the amplitude and spread of the excitatory signal, whereas 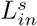 and 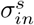 describe the inhibitory signal. 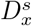 and 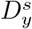 are the distances between the two neurons on both axes (computed as the differences between the RF centres), and the null term is used to avoid self-excitation. This equation is multiplied with the activation value of the pre-synaptic neuron in order to compute to post-synaptic lateral input.

### 2.3 Cross-Modal Connections

Furthermore, the modalities themselves are connected to each other via cross-modal synapses. This is motivated by the neurobiological findings of such connections (visuo-tactile: Blake et al. (2004), audio-visual: Molholm et al. (2002), audio-tactile: Foxe et al. (2000); for a survey see: Ghazanfar and Schroeder (2006)) These are modelled following the work of Cuppini et al. (2017). They opt for excitatory-only cross-modal connectivity, since there is poor evidence for such long-range cortico-cortical inhibitory synapses. Interestingly, Iurilli et al. (2012) observe rodent visual modality inhibition caused by links coming from the auditory modality, but it appears that those links would act as excitatory input for local inhibitory neurons, which end up causing GABAergic inhibition of the visual supragranular pyramids. Essentially, neurons from one sensory modality will exclusively excite neurons covering the same region of space in the other two sensory modalities. This is implemented through the following Gaussian function:

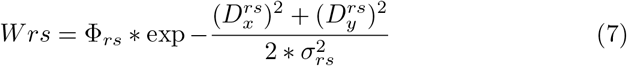

Where *r* and *s* represent the two regions that are being linked, Φ_*rs*_ the amplitude of the function and *σ*_*rs*_ the spread. The distance functions 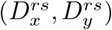 are analogues to the lateral connection distances, with the main difference being that the computation is between the RF centres of two neurons on differing modalities.

### 2.4 The Multisensory Neuron

The choice of a single multisensory neuron is motivated by findings of multi-modal neurons with RFs as large as the hand (Graziano et al., 1997; Rizzolatti et al., 1981). As in previous neurocomputational work, this model represents the entire system, not a single neuron.

The unimodal regions are all connected to the multisensory neuron through feedforward and feedback connections. Since this model is meant to simulate peri-hand plasticity, the visual and auditory region have their strongest links with the multisensory region in the limit of space covering the hand. Furthermore, the tactile feedback and feedforward synaptic weights were maintained uniform.

### 2.5 Feedforward Connections

The bottom-up connections from the unisensory region into the multisensory neuron are defined as follows:

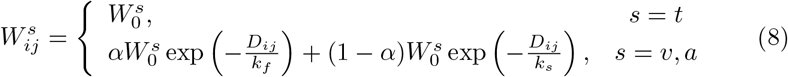

Here, 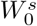 represents the synaptic strength within or adjacent to the spatial region encoding the hand. Parameter *α* determines the amplitude associated with each exponential component, whereas *kf* and *ks* denote the fast and slow decay rates, respectively. These parameters are calibrated to ensure an immediate and pronounced reduction in connectivity strength as the synapses traverse a peri-hand boundary (lim). The distances (*D*_*ij*_) are computed as:

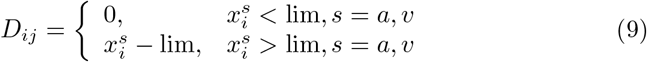

### 2.6 Feedback Connections

The feedback connections are similarly calculated:

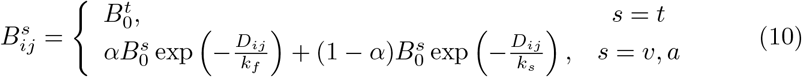

Here, each term maintains the same function as in the preceding equation, with the sole exception that the maximal value 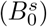 is incrementally larger, thereby simulating enhanced top-down connectivity (Siu et al., 2021; Douglas et al., 1995) (but see Choi et al. (2018)).

## 3 Network Activity

### 3.1 Inputs to the Unisensory Region

At every time step, the input to each neuron is computed as:

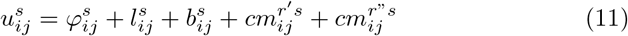

Here, 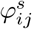 denotes the external input to the network, 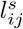 signifies the lateral input from proximal neurons within the same modality, 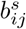 refers to the feed-back input from the multisensory neuron, and, ultimately, 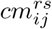 corresponds to the cross-modal input originating from neurons that encode analogous spatial regions in the other two modalities (thus represented by *r*′ and *r*′′). The subsequent equations delineate the aforementioned four components of the equation previously introduced:

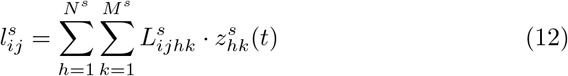

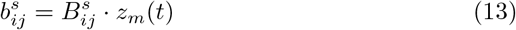

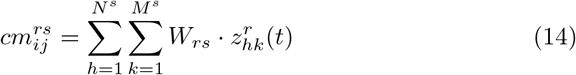

The expression 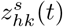 denotes the activity of the pre-synaptic neuron (*hk*) within the same modality at time step *t*. Likewise, *z*_*m*_(*t*) characterizes the activity of the multisensory neuron at time step *t*, whereas 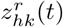 represents the activity of the pre-synaptic neuron (hk) from modality *r* at time step *t*.

### 3.2 Inputs to the Multisensory Region

The input to the multisensory neuron is derived from the aggregate convolution of the activity across individual unisensory regions with their corresponding feedforward weights:

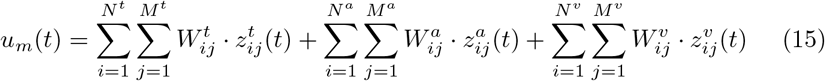

### 3.3 Steady-state Equations

The activity of a neuron is computed through a first order equation and a sigmoidal activation function:

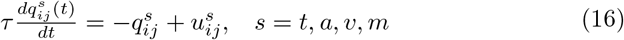

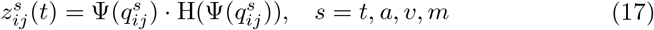

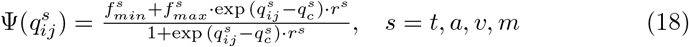

Here, 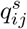 serves as the state variable, while 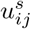 denotes the input function previously elaborated. 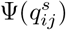 functions as the sigmoidal activation function. Notably, H(*·*) denotes the Hessian through which the sigmoidal is channelled, thereby ensuring non-negative neuronal activity. Moreover, *τ* symbolizes the time constant of the differential equation, corresponding to the membrane time constant. Within the parameters governing the sigmoidal function, 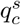 designates the central point, *r*^*s*^ specifies the slope, and 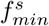 and 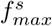 indicate the lower and upper boundaries, respectively.

### 3.4 Input

In the experimental setup the tactile input was an electrical current applied to the finger. We modelled the tactile input as being delivered at position 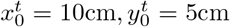 of the tactile area. The auditory input were approaching sounds at different velocities. We modelled the sound stimuli as being delivered at varying distances from the individual, represented by position 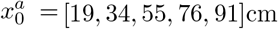 in the auditory area, while maintaining a constant position in the complementary axis at 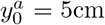. The input stimulus to the network is depicted as a Gaussian blob:

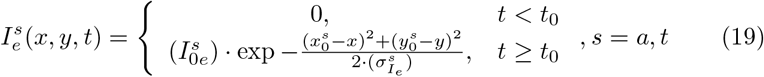

The amplitude of the function 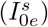 denotes the stimulus intensity, whereas the spread 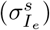 signifies the spatial extent. As previously noted, the experimental procedure is conducted with the participant blindfolded, ensuring the exclusion of any visual input.

## 4 Training simulation in the network

To recreate the active tool use training in the model, we do not explicitly model motor processing and motor neurons activity but only the sensory consequences of each object movement as a synchronous audio-visuo-tactile stimulus in the network. Following Serino et al. (2015), we assume that the synchronicity between tactile stimulation in the hand and auditory or visual stimulation in the far space triggers the expansion of PPS via Hebbian Learning.

The audio-visuo-tactile stimulus, *I*^*s*^ (*x*^*s*^, *y*^*s*^), used to represent the sensory consequences of object movements is described by the following equation:

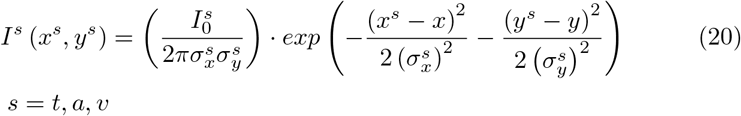

Here, 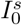 represents the intensity of the stimulus, while 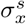 and 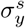 represent its vertical and horizontal spread, respectively (where s is either tactile, auditory or visual). The tactile stimulus is always delivered at the central point of the tactile area, whereas auditory and visual stimulus locations are always centred on the vertical axis.

In line with Serino et al. (2015), we modelled network plasticity following a Hebbian Learning rule for auditory and visual feedforward synapses. The change in the synaptic weight, 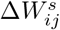, at each time step during the training simulation is defined as:

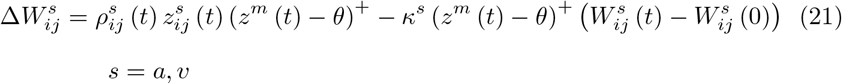

Here, the subscript indices *i* and *j* denote the specific location of the presynaptic neuron along the horizontal and vertical axes of the unisensory area *s*, respectively. The first term of the equation represents the reinforcing factor, with the activities of the pre-synaptic 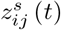 and multisensory post-synaptic neurons *z*^*m*^ (*t*) multiplied by the learning rate 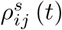 (where *s* is either auditory or visual^1^). This rate adopts a starting value 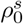 and gradually reduces as the connection strengthens, reaching zero when the weight is at its maximum, while the multisensory neuron is weighted against a small threshold *θ* to avoid strengthening in cases of residual activity. The second term of the equation is the forgetting factor, which is computed when the pre-synaptic neurons are inactive and the post-synaptic neuron has above-threshold activity. Here, the feedforward synaptic weights 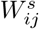 are penalised by the forgetting rate *κ*^*s*^.

For simplicity, instead of modelling the sensory consequences of 100 movements, the audio-visuo-tactile stimulus was presented 10 times to train the network. Each stimulus lasted 500 simulation steps (i.e., 200 ms). To reproduce the sensory effects of motor training in the network, we fitted the parameters that define the network learning rate 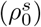, the horizontal position (*y*^*s*^) and the spread 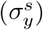 of audiovisual stimuli, ensuring that post-training the HC model RTs match the post-training PPS representation of the HC group (Ferroni et al., 2022).

## 5 Modelling the influence of SCZ in the network

### 5.1 Modulation of the E/I balance

The modulation of the E/I balance was implemented by increasing or decreasing the strength of the recurrent excitatory connectivity in each sensory area. This was done by modifying the value of the parameter 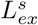, defined in Equation 22:

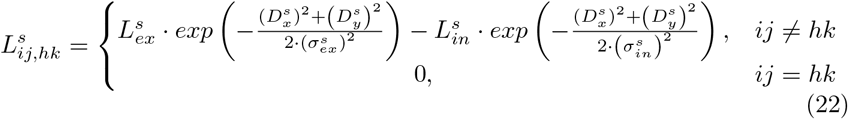

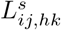 denotes the weight of the synapse from the pre-synaptic neuron at position *hk* to post-synaptic neuron at position *ij* ^2^. 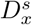 and 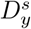 indicate the distances between the pre-synaptic neuron and the post-synaptic neurons along the horizontal and vertical axis of the unisensory area. The excitatory Gaussian function is defined by the parameters 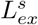 and 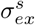, whereas the inhibitory function is defined by 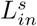 and 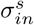. The model assumes no auto-excitation (i.e. 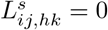, if *ij* = *hk*). The change in the E/I ratio was modelled by increasing or decreasing the value of the parameter 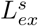 in all unisensory areas.

### 5.2 Failures in top-down signalling

The failures in top-down signalling were implemented by uniformly weakening top-down synaptic weights between multisensory and unisensory areas. This was achieved by changing the value of the parameter 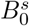, defined in Equations 23 and 24:

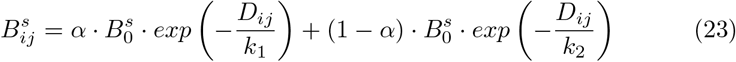

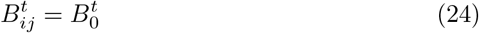

Here, the distance *D*_*ij*_ is equal to zero for the auditory neurons that encode the first *Lim* cm of the auditory space, while for the neurons outside this boundary *D*_*ij*_ is the minimum Euclidean distance between its RF centre and this boundary. 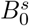 denote the value of the feedback and feedforward synapses, respectively, when *D*_*ij*_ is equal to zero in the auditory and visual areas. *k*_1_, *k*_2_ and *α* are parameters governing the exponential decay of the synaptic weights of auditory neurons encoding regions outside of the near-space of the hand.

## 6 Fitting

### 6.1 Sigmoid fitting of network output

The RTs values generated by the network in the simulation of the experiment were fitted to a sigmoid function described by Equation 25.

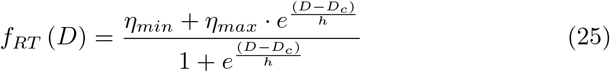

The term *D* denotes the distance from the hand in cm at which the auditory stimulus was applied. The parameters *η*_*min*_ and *η*_*max*_ represent the lower and upper saturation of the sigmoidal relationship. *D*_*c*_ is the central point of the function and *h* denotes the slope of the function at its central point.

Parameters *D*_*c*_ and *h* were estimated by a least-squares fitting procedure using the Trust Region Reflective algorithm available in the SciPy library for the Python programming language (Virtanen et al., 2020). The central point was bounded to be positive, whereas the slope was unbounded. The initial guesses given to the algorithm were defined according to equations 26 and 27.

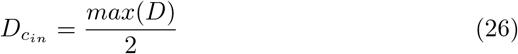

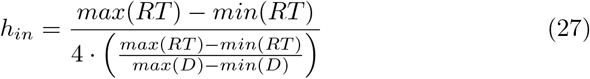

Here, 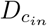 and *h*_*in*_ denote the initial guesses of the central point and the slope respectively. The arguments *max* and *min* refer to the maximum and minimum value of a given vector respectively. The estimated value of the sigmoid central point (*D*_*c*_) was assumed as the boundary of the PPS representation generated by the model and the value of the slope (*h*) the sharpness of its definition.

#### 6.1.1 Sigmoid fit or Spearman-Karber method

The empirical data (Ferroni et al., 2022) modelled in this study show different slope results before and after training depending on the method used to calculate the psychometric curves: sigmoid fit or Spearman–Karber method. Presumably, this discrepancy occurs due to the few time points (i.e. 5 delays from 300 to 2700 ms) measured in the experiment. This discrepancy is problematic for our modelling results, given that the slope of the PPS representation generated by our model is very sensitive to the learning rate of the network (see Figure 3).

We used the sigmoid fit procedure instead of the Spearman-Karber (SK) method because the network’s mathematical model produces a sigmoidal response shape. Hence, our model presumes a sigmoidal psychometric function for the simulated experiment which the SK method does not. Using a sigmoidal psychometric function affects the data used for model fitting, since achieving a sigmoid fit implies excluding 17 participants (6 in the HC group and 11 in the SCZ group) due to poor sigmoidal fit (*R*^2^ < .25).

Notably, the average RT per group without excluding participants forms a peak, not a sigmoid shape (see Supplementary Figure 2 in Ferroni et al. (2022)). In contrast, our network model assumes that RTs rise exponentially after the close space of the hand, forming a sigmoid-like shape. This RT change stems from the exponential decrease of feedforward/feedback synaptic weights outside the near-space of the hand. This core assumption of our model must remain unchanged; otherwise, a new PPS network model would be needed, which is beyond the scope of this study.

### 6.2 Fitting to RT data of participants

We replicated the procedure described in Paredes et al. (2022) to match the RT generated by the model to the RT observed in the experiment participants. This procedure consisted in matching the time units of the model and the experiment and minimising a cost function with a stochastic optimisation algorithm. At each iteration of the minimisation routine, the RTs were matched by a standard linear regression, as presented in Equation 28.

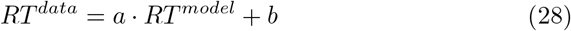

Here, *RT*^*data*^ and *RT*^*model*^ represent the RTs obtained in the empirical study and the experiment simulation respectively. Parameter *a* denotes how many milliseconds correspond to one time unit of the network model, while parameter *b* represents the duration of neural processing not captured by the model.

During the optimisation, negative values of *a* and *b* were converted to 0 because negative coefficients do not make sense according to this definition. The employed cost function is defined by Equation 29.

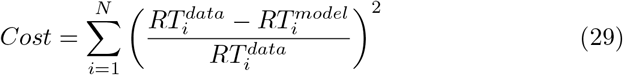

Here, 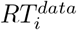 and 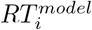 denote the RT measured at the *i* th distance point. N represents the number of distances measured (e.g. 5 in the empirical study).

This cost function was minimised by the implementation of the differential evolution algorithm available in the SciPy library for the Python programming language (Virtanen et al., 2020).

For the fitting, we considered 15 equally spaced distance points in the range between 19 and 91 cm. *RT*^*model*^ is composed of 15 RTs generated by the model at those distance points.

*RT*^*data*^ is composed of 15 RT taken from a sigmoid curve representing the group-level response profile of each group for each condition. This curve was generated from the median central point and median slope of each group. The minimum and maximum values of the curve were calculated from the median RT of the group. The parameters generating the sigmoid curve for each group are presented in Table 1.

**Table 1:** Parameters of the sigmoid curve representing the data of each group before and after tool use.

|  | CP | Slope | Min | Max |
| --- | --- | --- | --- | --- |
| HC pre-training | 56.56 cm | .043 | 270 ms | 318.5 ms |
| HC post-training | 72.40 cm | .085 | 271.5 ms | 326.5 ms |
| SCZ pre-training | 50.76 cm | .045 | 382.5 ms | 429.5 ms |
| SCZ post-training | 64.72 cm | .062 | 373 ms | 455 ms |

To select the models that provide the best fit to the data, we employed a simple model comparison approach based on the Root Mean Square Error (RMSE) adjusted by the number of free parameters, as defined in equation 30:

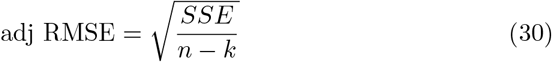

Here, SSE stands for Sum of Squared Error, n denotes the sample size, and k refers to the number of free parameters.

## 7 Accounting for PPS differences in SCZ before training

We systematically varied the parameters governing recurrent excitation 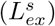, feedforward pruning (*ρ*_*W*_ ^*s*^), top-down weights 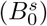 and cross-modal pruning (*ρ*_*W*_^*ss*^) in the pre-training HC model to explore plausible mechanisms behind the differences in PPS observed in SCZ before training (see Supplementary Figure 1). Our simulations revealed that an increase in recurrent excitation and top-down synaptic weights reduces the size of the PPS representation generated by the model. In contrast, the decrease in synaptic density in auditory feedforward connections influences both the size and the slope of the PPS. No relevant effects were observed for cross-modal pruning. None of the explored parameters result in a shallower slope, so our model never matches the non significant slope reduction seen in the SCZ group before training (Figure 1B). Despite the aforementioned modifications to the network and the experimental data being modelled, the influence of the model parameters on the size and slope of the PPS representation remains consistent with Paredes et al. (2022).

**Figure 1:**
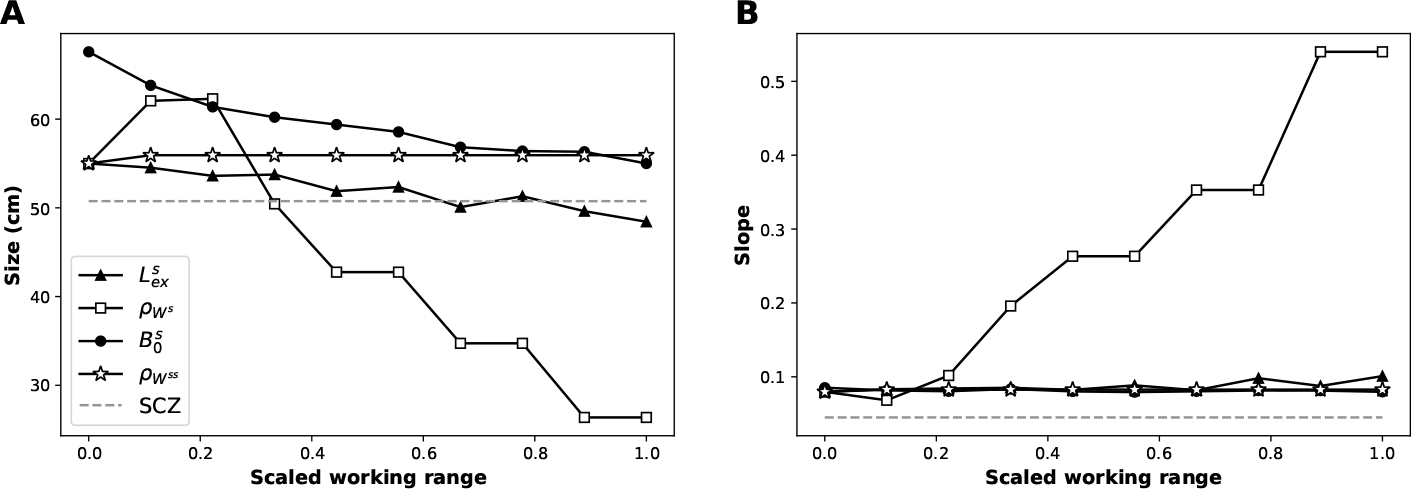
Effects of alterations in neural mechanisms on the size and slope of the PPS representation generated by the HC pre-training model. **Panels A** and **B** show the effect of systematic variation of parameters in the range at which they produce sigmoid-like PPS representations (these ranges were scaled to facilitate visual comparison). The dashed lines indicate the size and slope observed reported in the experimental data (Section 2). The size reduction observed in SCZ could be reproduced by either an increase in lateral excitation 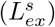 or pruning of feedforward synapses 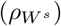.

## 8 Reproducing PPS expansion in HC with the SCZ model

Our modelling results are built on top of the assumption that all numerical differences in the parameters that define the sigmoid curves that represent the PPS representation between groups and conditions are relevant (Table 1). This data shows that PPS expansion in SCZ is smaller than in Controls. Accounting for this difference in our model fitting can be perceived as problematic, as statistical analysis shows that these differences are not statistically significant. As an additional verification analysis, we reran our simulations using, as posttraining data for the SCZ group, a sigmoid function constructed by assuming that the PPS extension is actually identical in controls and SCZ. The parameters generating the sigmoid curves in this new set of simulations are presented in Table 2.

**Table 2:** Parameters of the sigmoid curve representing the data of each group before and after tool use in this new set of simulations. In bold are the parameters that were changed compared to the ones that were used for the simulations reported in the article.

|  | CP | Slope | Min | Max |
| --- | --- | --- | --- | --- |
| HC pre-training | 56.56 cm | .043 | 270 ms | 318.5 ms |
| HC post-training | 72.40 cm | .085 | 271.5 ms | 326.5 ms |
| SCZ pre-training | 50.76 cm | .045 | 382.5 ms | 429.5 ms |
| **SCZ post-training** | **66.58 cm** | **.087** | **384 ms** | **437.5 ms** |

We observe that we still need to reduce synaptic plasticity in the network to improve the fit to the post-training data (see Figure 2). However, now the improvement in goodness of fit is not as great as in the simulations considering the actual SCZ post-training data (2.88 ms vs 0.66 ms).

As before, we achieved an equivalent fit to the new SCZ post-training data after reducing the learning rate (from 6.14 × 10^−2^ to 3.84 × 10^−2^), increasing the forgetting rate (from 5 × 10^−5^ to 4.20 × 10^−4^), or increasing the plasticity threshold (from .05 to .44). The results of these possible models are shown in Table 3.

**Table 3:**
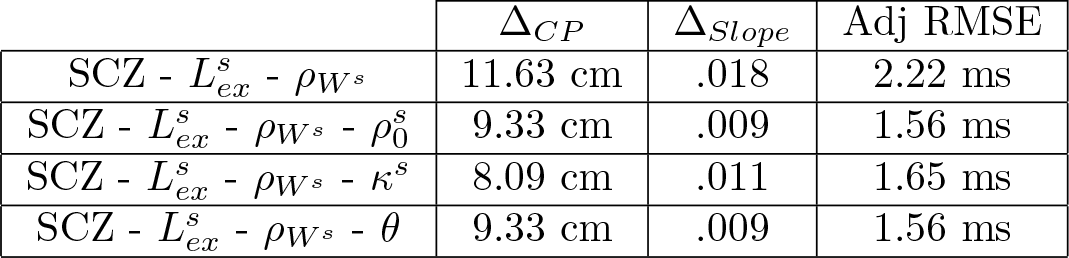
Effects of tool use on PPS representation observed in the evaluated models. Δ represents the difference between post-training and pre-training. Adj RMSE are calculated against the sigmoid curve generated by the SCZ pretraining data shown in Table 2.

**Figure 2:**
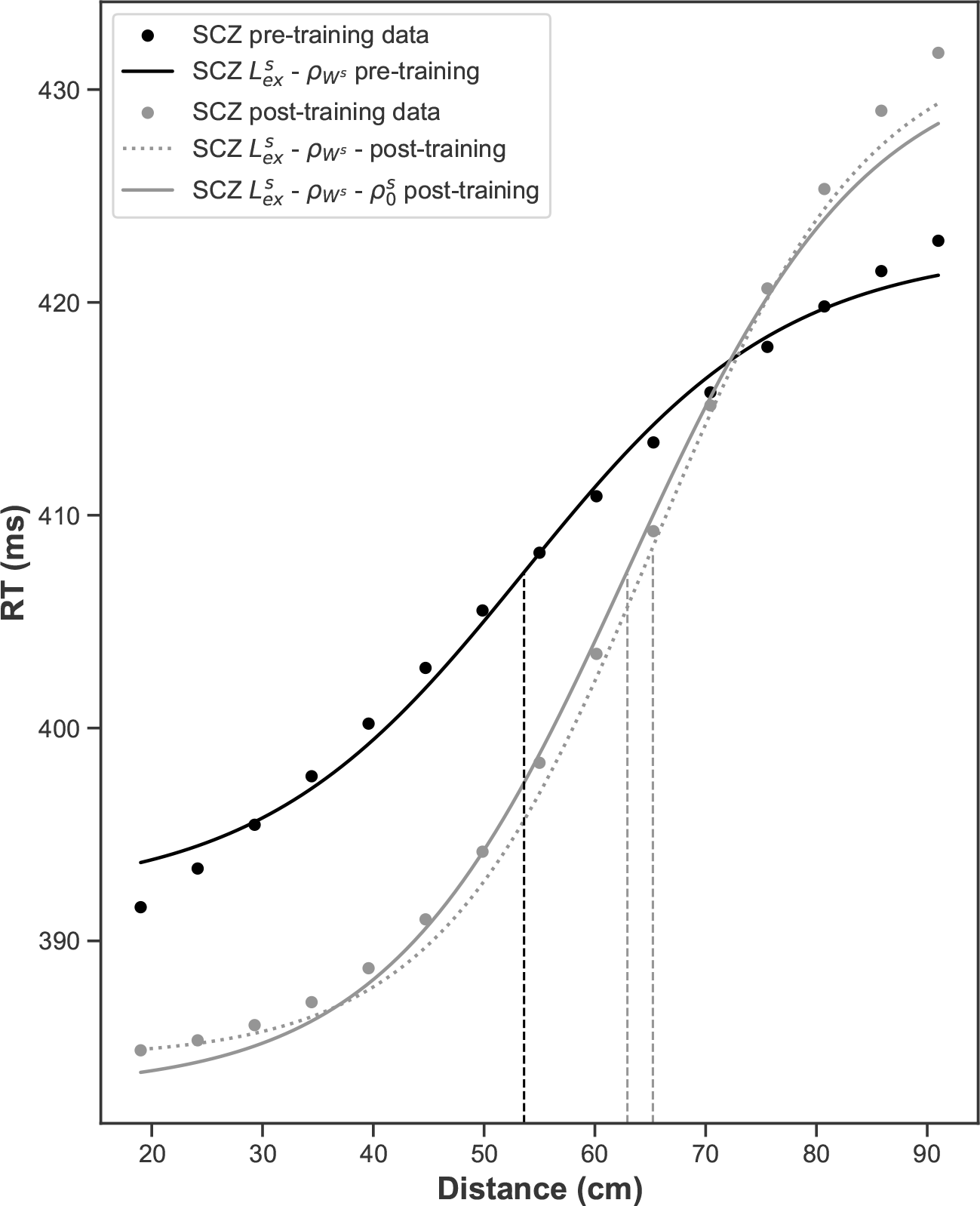
PPS representations generated by the SCZ network models. The solid lines depict the sigmoid curves characterising the data generated by the models. The dots represent the data collected in the experiment (Section **??**). Vertical dashed lines mark the central points of the sigmoid curves, interpreted as the sizes of the PPS representations. Increased recurrent excitation 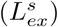 or pruning of feedforward synapses 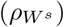 are required to generate a close match to the SCZ pre-training data (black dots). The grey dotted line depicts the result of applying the simulated training routine to the SCZ pre-training model. The grey solid shows how a model with a reduced learning rate 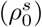 achieves a good fit to the SCZ post-training data (grey dots).

In contrast, in this new set of simulations, we observe differences in synaptic weight increase after training between the possible models (see Figure 3).

**Figure 3:**
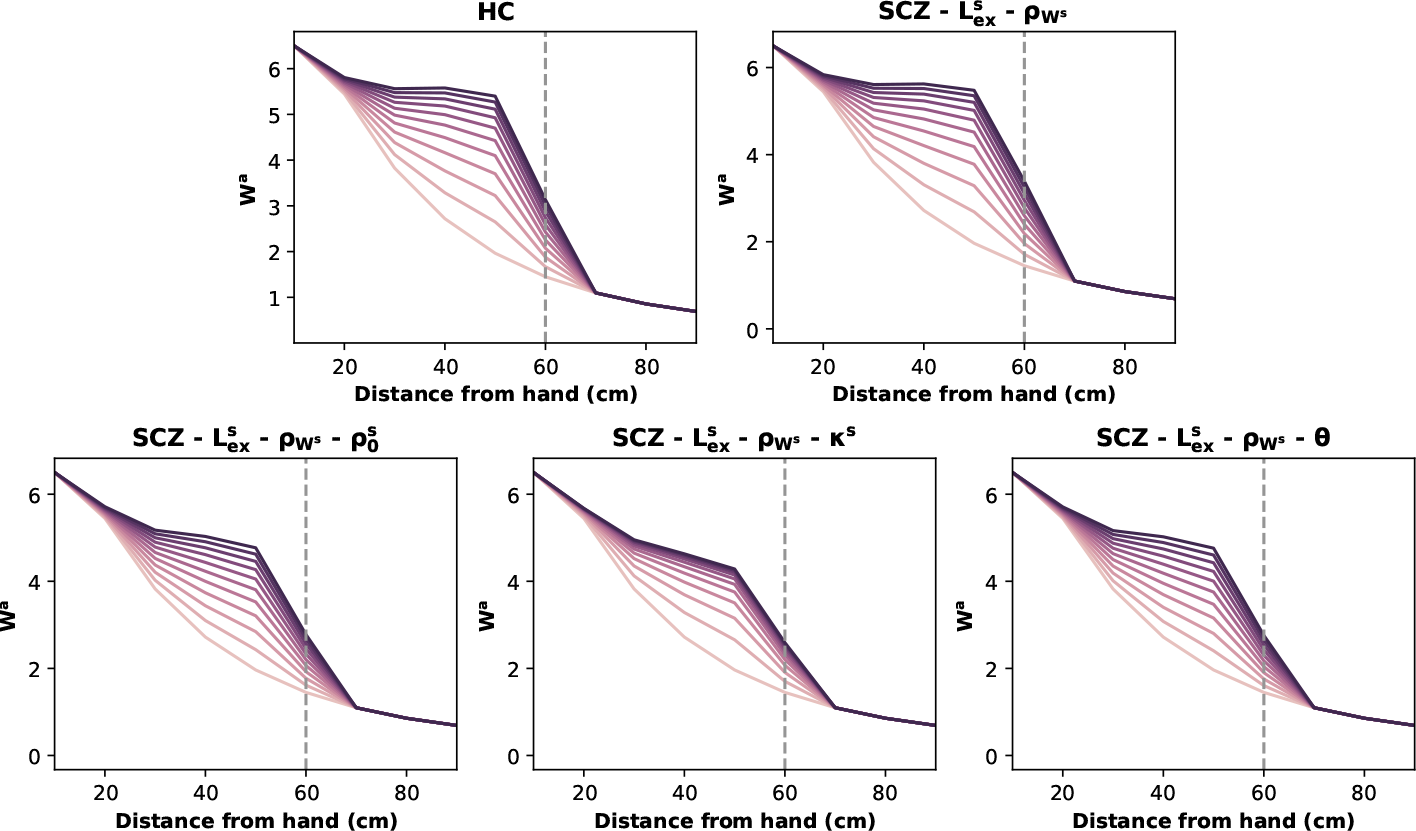
Effects of training in feedforward synaptic weights in the examined network models. The plots show the feedforward synaptic weights of the neurons encoding the auditory space (*W*^*a*^) as a function of the distance from the hand (*N*^*a*^ = 2). The gray dashed line represents the distance at which the audiovisual stimulus was presented. The colour-coded lines depict the evolution of the weights during a 10 step training, with lighter shades at the beginning and darker at the end. Compared to baseline HC and SCZ models, weakening the reinforcing factor 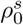 or *θ*) leads to weaker synaptic learning, whereas strengthening the forgetting factor (*κ*^*s*^) leads to an even weaker synaptic learning that converges in fewer steps.

Overall, this verification analysis shows that our results hold even after assuming the exact same PPS plasticity between HC and SCZ groups. However, the observed reduction in network plasticity is subtler.

## 9 Predictions

### 9.1 PPS expansion after long training sessions

We simulated the training routine with the same parameters as in Figure 4, increasing the training steps (stimulus presentations) from 10 to 60. This longer stimulation allows us to distinguish more clearly the outcomes of the competing mechanistic accounts (see Supplementary Figure 4). We observe that now the three models do not converge to the same solution. The increased forgetting rate model (*κ*^*s*^) does not increase auditory synaptic weights (*W*^*a*^) as much as the reduced learning rate model 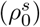 or the increased plasticity threshold model (*θ*). The resulting effects on the size and slope of the PPS representations are shown in Table 4.

**Figure 4:**
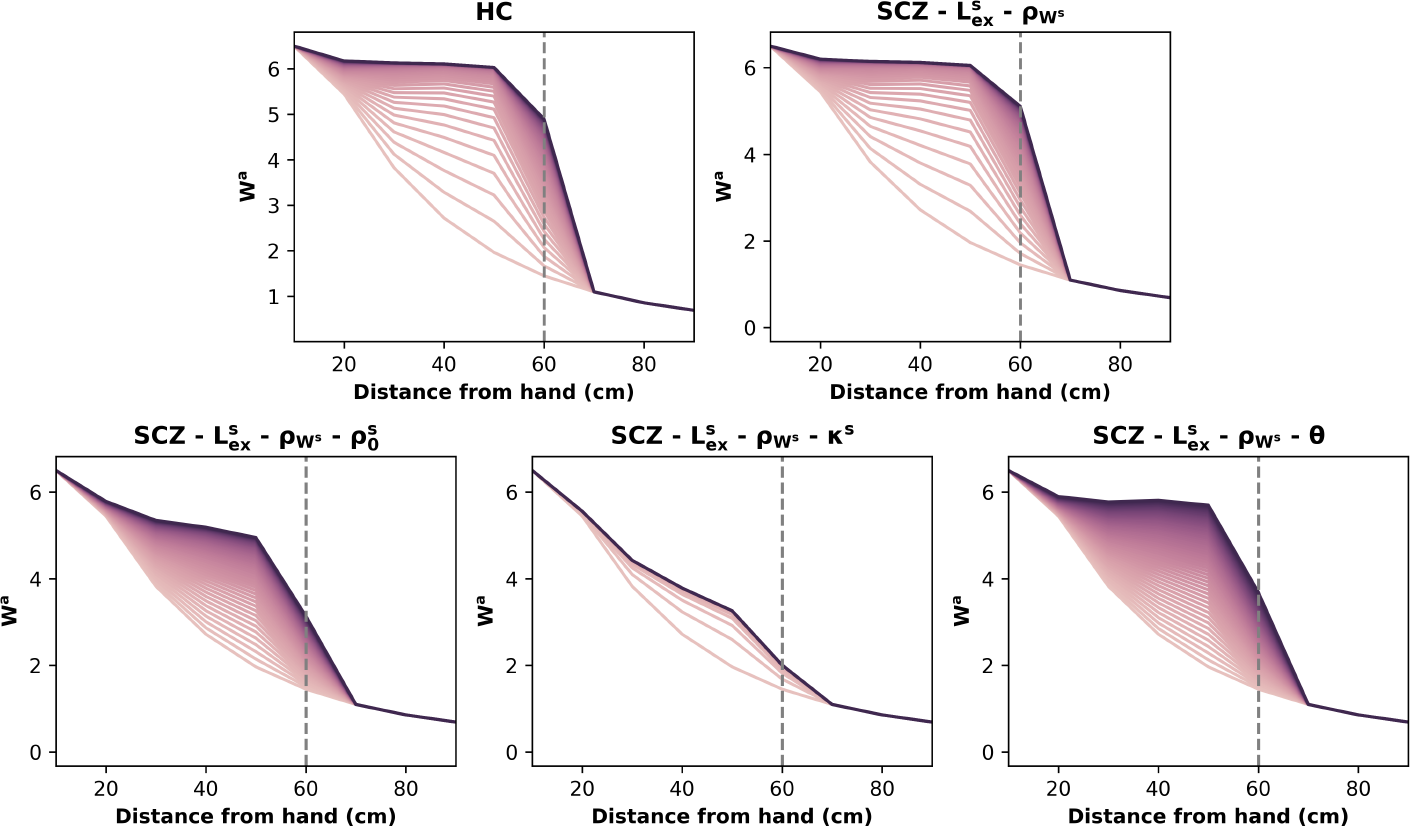
Effects of a longer training session in feedforward synaptic weights in the examined network models. The plots show the feedforward synaptic weights of the neurons encoding the auditory space (*W*^*a*^) as a function of the distance from the hand (*N*^*a*^ = 2). The gray dashed line represents the distance at which the audiovisual stimulus was presented. The colour-coded lines depict the evolution of the weights during training, with lighter shades at the beginning and darker at the end.

**Table 4:** Effects of long training session on PPS representation observed in HC and SCZ models. Δ represents the difference between post-training and pretraining.

| | $\Delta_{CP}$ | $\Delta_{Slope}$ |
| --- | --- | --- |
| HC Model | 14.77 cm | .051 |
| SCZ - $L_{ex}^s - \rho_{W^s}$ | 15.18 cm | .037 |
| SCZ - $L_{ex}^s - \rho_{W^s} - \rho_0^s$ | 10.57 cm | .016 |
| SCZ - $L_{ex}^s - \rho_{W^s} - \kappa^s$ | 4.45 cm | .001 |
| SCZ - $L_{ex}^s - \rho_{W^s} - \theta$ | 12.25 cm | .024 |

### 9.2 PPS expansion after training at a shorter distance

We simulated the training routine with the same parameters as in Supplementary Figure 2, reducing the distance at which the stimulus was presented from 60 to 30 cm. Here we observe that the three models also do not converge to the same solution. The increased forgetting rate model (*κ*^*s*^) does not increase auditory synaptic weights (*W* ^*a*^) as much as the reduced learning rate model 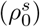 or the increased plasticity threshold model (*θ*). The resulting effects on the size and slope of the PPS representations are shown in Table 5.

**Figure 5:**
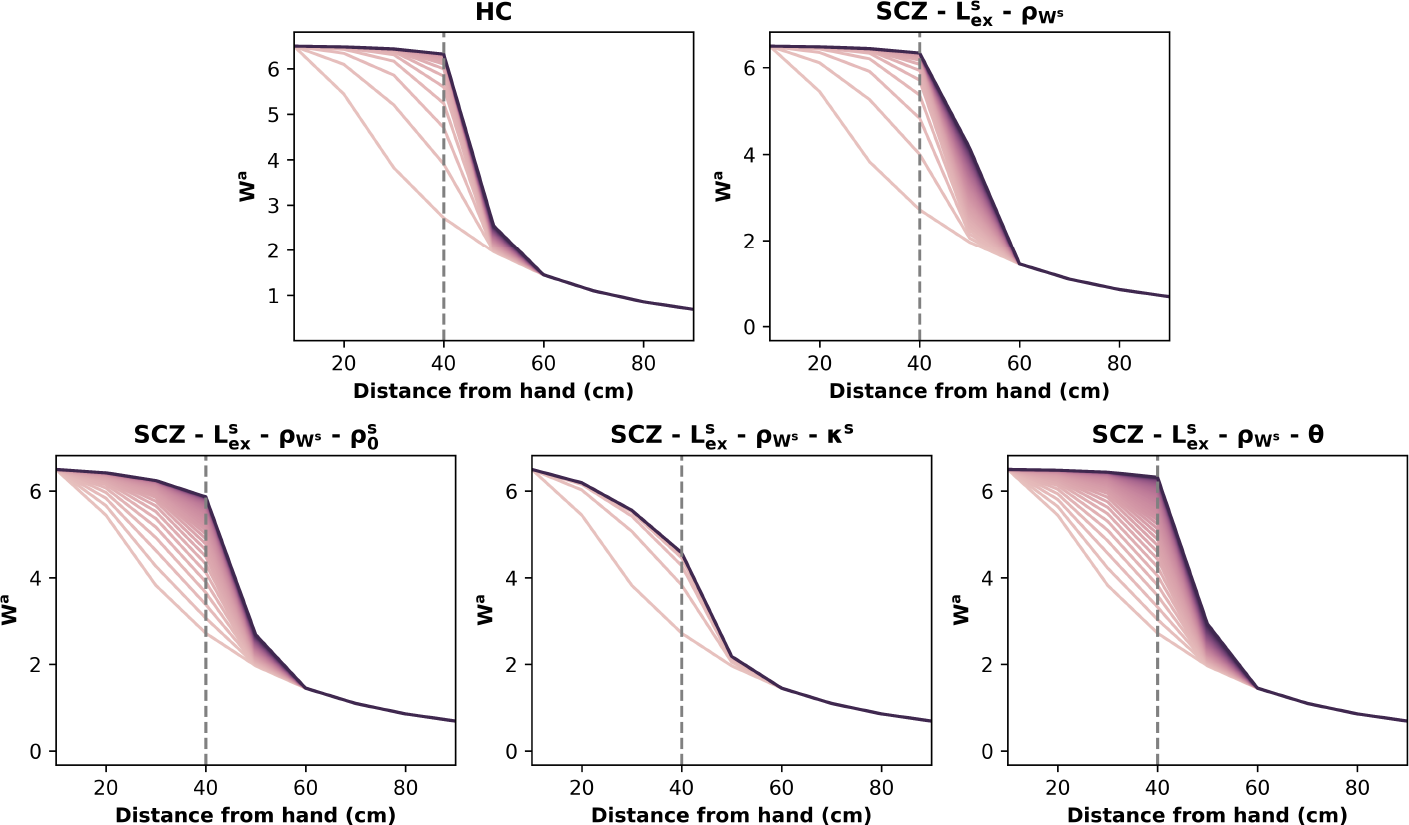
Effects of a long training session at a shorter distance in feedforward synaptic weights in the examined network models. The plots show the feedforward synaptic weights of the neurons encoding the auditory space (*W*^*a*^) as a function of the distance from the hand (*N*^*a*^ = 2). The gray dashed line represents the distance at which the audiovisual stimulus was presented. The colour-coded lines depict the evolution of the weights during training, with lighter shades at the beginning and darker at the end.

**Table 5:**
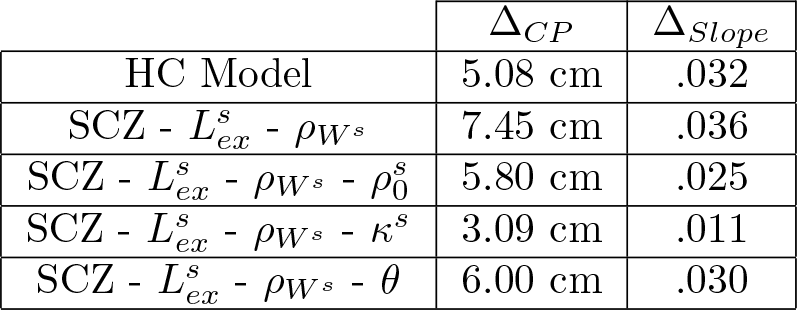
Effects of long training session with stimulus at a shorter distance on PPS representation observed in HC and SCZ models. Δ represents the difference between post-training and pre-training.

| | $\Delta_{CP}$ | $\Delta_{Slope}$ |
| --- | --- | --- |
| HC Model | 5.08 cm | .032 |
| SCZ - $L_{ex}^s$ - $\rho_{W^s}$ | 7.45 cm | .036 |
| SCZ - $L_{ex}^s$ - $\rho_{W^s}$ - $\rho_0^s$ | 5.80 cm | .025 |
| SCZ - $L_{ex}^s$ - $\rho_{W^s}$ - $\kappa^s$ | 3.09 cm | .011 |
| SCZ - $L_{ex}^s$ - $\rho_{W^s}$ - $\theta$ | 6.00 cm | .030 |

## 10 Parameter values of the HC model

**Table 6:**
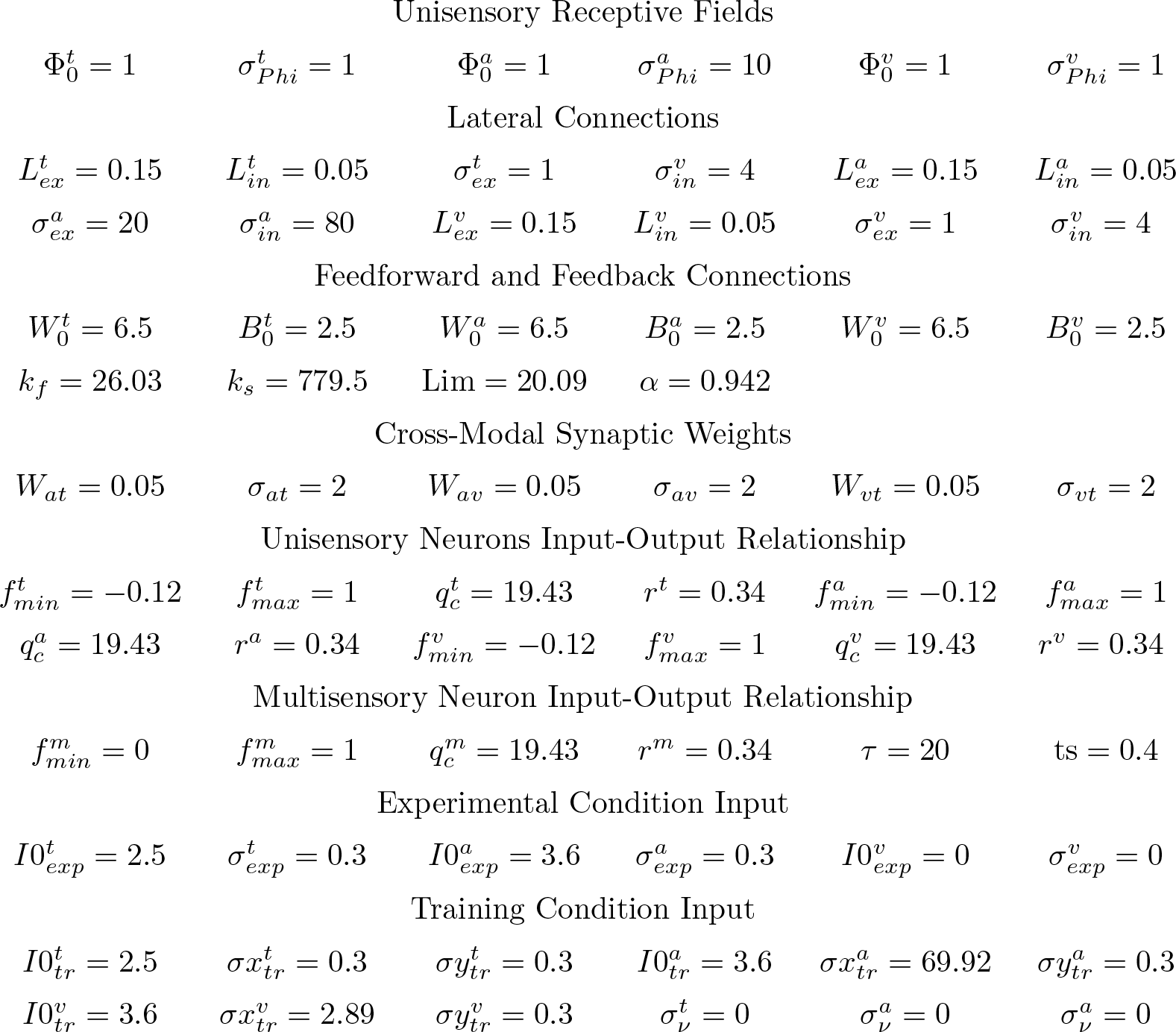
Parameter values in the HC model. This parameterisation was kept fixed throughout the simulations calculated to account for the SCZ model.

## Footnotes

1 Tactile neuron synaptic weights are constant and not dependent on body distance, so they are excluded from plasticity computations.

2 Postmortem brain studies in SCZ exhibit considerable heterogeneity, with no consistent evidence of reduced synaptic density within the cortical regions being modelled.

1 Tactile neuron synaptic weights are constant and not dependent on body distance, so they are excluded from plasticity computations.

2 Subscripts *i* and *j* indicate neuron positions in unisensory area *s*, with s as either *t* (tactile), *v* (visual), or *a* (auditory).

## Notes

### Competing Interest Statement

The authors have declared no competing interest.

### Summary of Updates

Overall writting updated to clarify. All Figures were revised to simplify methodological and results report.

